# Neutralizing PD-L1 and PD-L2 Enhances the Efficacy of Immune Checkpoint Inhibitors in Ovarian Cancer

**DOI:** 10.1101/2020.01.19.911941

**Authors:** Yu Rebecca Miao, Kaushik N. Thakkar, Jin Qian, Mihalis S. Kariolis, Huang Wei, Saravanan Nandagopal, Teddy Yang, Anh N. Diep, Gerald Maxwell Cherf, Yu Xu, Eui Jung Moon, Yiren Xiao, Haizea Alemany, Tianer Li, Wenhua Yu, Bo Wei, Erinn B. Rankin, Amato J. Giaccia

**Author notes:** To whom correspondence should be addressed: Amato J. Giaccia.

## Abstract

Immune checkpoint inhibitors targeting the PD-1/PD-L1 pathway have improved for a number of solid tumors. Unfortunately, ovarian cancer represents a major clinical hurdle for immune checkpoint blockade (ICB) with reported low patient response rates. Using IHC staining, we find that PD-L2 is highly expressed in ovarian cancers and other malignancies with sub-optimal response to ICB, and is expressed at low levels in cancers responsive to ICB. Based on this observation, we hypothesized that the elevated expression of PD-L2 produced by both tumor and surrounding stromal cells contributes to immune-suppression. Since PD-L2 has been reported to have a 6- to10-fold higher native binding affinity to PD-1 compared with PD-L1, we hypothesized that high levels of PD-L2 can lead to insufficient blockade of the PD-1 signaling pathway. To overcome the immune repressive activity of PD-L2, we engineered a soluble PD-1 decoy molecule (sPD-1 mutant) that binds and neutralizes both PD-L1 and PD-L2 with a 10,000- and 200- fold improvement in binding affinity, respectively, when compared to wild-type binding to these same molecules. Such enhancement in binding affinity is facilitated by amino acid mutations both within and outside of the binding interface. Furthermore, this high affinity sPD-1 mutant molecule demonstrates superior *in vivo* efficacy in multiple cancer models including ovarian cancer where PD-L2 is highly expressed on the cell surface.

**One Sentence Summary:** Dual Inhibition of PD-L1 and PD-L2 using an affinity enhanced sPD-1 decoy molecule delivers superior antitumor activity when compared with αPD-1 and αPD-L1 antibodies in ovarian cancer.

## Introduction

Therapeutic inhibition of the PD-1 signaling pathway has proven to be an effective strategy in treating malignant diseases, showing unprecedented long-term durable responses in the clinic (1–3). During the last decade, therapeutic antibodies against PD-1 and PD-L1 were approved for the treatment of various cancer types either as monotherapies or in combination with current standard-of-care therapeutic regimens (4, 5). However, clinical responses can vary significantly between patients and reflect the heterogeneous nature of individual tumors and their microenvironment (TME) (4, 6, 7). Ovarian cancer is known to respond poorly to immune checkpoint blockade (ICB) with clinical trials reporting responses ranging from 6%-22% (8, 9). Clearly, there is a knowledge gap between our current understanding of immune regulation and how it can be manipulated to enhance the clinical efficacy of ICB therapies in ovarian cancer. An ongoing effort to enhance the response rate of ICB inhibitors is being pursued in the clinic through combining these inhibitors with cytotoxic drugs, targeted therapies and radiation treatment (10–12). However, responses to ICB in ovarian cancer are still low, even in combination with cytotoxic chemotherapy and targeted therapy (9). These clinical results provide an explanation for why there has yet to be a FDA approved ICB therapy for ovarian cancer.

In contrast to the low response to ICB therapy, high-grade serous ovarian cancer was one of the first human cancers to have demonstrated a relationship between increased tumor-infiltrating lymphocytes (TILs) and increased patient survival (8, 13). A major challenge remains to boost the clinical efficacy of ICB in ovarian cancer and overcome low mutational burden and an immunosuppressive tumor microenvironment comprised of regulatory T cells, macrophages and PD-1 signaling (14) . Therefore, strategies to overcome the immunosuppressive TME through complete inhibition of the PD-1 signaling pathway should facilitate enhanced TIL infiltration and activity, and enhance tumor control of ovarian cancer.

Mechanistically, most studies have exclusively focused on the role of PD-L1 on PD-1 signaling and have failed to explore the consequences of PD-L2 activating PD-1 signaling in cancer. A second ligand to PD-1, PD-L2 is generally expressed at low levels on dendritic cells, macrophages and endothelial cells during physiological homeostasis (15, 16). However, stimulation by pro-inflammatory cytokines such as IL-4 can induce PD-L2 expression on both immune and stromal cells (17, 18). Additionally, NF-kB and STAT6 signaling can also regulate PD-L2 expression on immune infiltrates (19).

Structural analysis of PD-1/PD-L1 and murine PD-1/PD-L2 show differences in binding modality. Binding of PD-L1 to PD-1 involves conformational change; whereas the binding of PD-L2 to PD-1 occurs in a direct manner (20). This difference in binding modality between PD-L1 and PDL-2 with PD-1 can explain the reported 6-10 fold increase in binding affinity between PD-L2 and PD-1 compared to PD-L1 and PD-1 (21). If both ligands are present at similar expression levels, PD-L2 would be predicted to outcompete PD-L1 binding to PD-1. Due to this binding advantage, high PD-L2 would also have a competitive advantage over therapeutic antibodies blocking the PD-1 receptor.

In this study, we investigated the expression of PD-L1 and PD-L2 in ovarian cancer stratified according to tumor grade. To overcome the problem of high affinity PD-L2 binding to PD-1, we developed a soluble PD-1 ligand trap (sPD-1) with an enhanced binding affinity to both PD-L1 and PD-L2. Our mutant sPD-1 has a higher apparent binding affinity for PD-L1 (10,000-fold) and PD-L2 (200-fold) when compared to wild-type PD-1 interactions with these ligands. Computational analysis of mutations identified in sPD-1 through yeast surface display screening reveals that enhanced binding is the consequence of amino acid changes in both the binding interface well as in non-binding interface of sPD-1. This enhancement in binding affinity translated into improved inhibition of PD-1 receptor activity both *in vitro* and *in vivo* models, outperforming antibodies targeting PD-L1 and PD-1 in two syngeneic ovarian cancer models and other cancer types. Our studies also indicate that PD-L2 alone can be an independent contributor towards immune escape. Therefore, the use of high-affinity sPD-1 mutant is a clinically viable strategy for overcoming the immune-suppressive TME present in ovarian cancer.

## Results

### PD-L2 is abundantly expressed in human ovarian cancers but not in ICB sensitive bladder cancer

Although PD-L1 and PD-1 interaction has been the subject of extensive study, less is known about the role of PD-L2 and PD-1, particularly in cancer types that respond poorly to anti-PD-1/PD-L1 therapies (22, 23). To elucidate the clinical relevance of PD-L2 expression in ovarian cancer, historically known to have a sub-optimal clinical response to checkpoint inhibitors, we analyzed PD-L1 and PD-L2 expression in human ovarian cancer tissue microarrays (n=156) (Fig. 1A). The specificity of anti-PD-L1 and anti-PD-L2 antibodies was validated in human tonsillar tissue (Sup. Fig.1A). The spatial distribution and staining intensity of PD-L1 and PD-L2 expression were quantified and stratified according to tumor grades. In ovarian cancer patients, the expression of both PD-L2 and PD-L1 are significantly elevated in the cancerous tissue compared to non-malignant specimens but did not appear to change significantly with tumor grade (Fig. 1B, 1C). Pearson correlation analysis showed a positive correlation between PD-L1 and PD-L2 expression in ovarian cancer, however non-overlapping expression pattern between PD-L1 and PD-L2 in ovarian cancer suggested that the two ligands may have different spatial expression (Sup. Fig. 1B). To evaluate whether high PD-L2 expression is a potential biomarker for low ICB responding cancer types, we also examined PD-L1 and PD-L2 expression in gastric cancer (n=76), esophageal cancer (n=72) and glioblastoma (n=152) tissue microarrays (TMA). These cancers also reported as having poor clinical respond to ICB inhibitors (24–26). Similarly, high PD-L2 expression was observed in tumors samples but not normal tissues (Sup. Fig. 1C-E). In contrast, bladder cancers have a reported higher clinical response rate to ICB (27) and interestingly, the staining of bladder cancer TMAs show low PD-L2 expression, but predominant PD-L1 ligand expression in bladder cancer cells (n=208) (Fig. 1D and 1E). In summary, we found cancers that are poor responders to ICB express high levels of both PD-L1 and PD-L2, whereas cancers that are sensitive to ICB almost exclusively express PD-L1. Based on this observation, we hypothesize this resistance to ICB is in part attributed to elevated PD-L1 and PD-L2 expression. Therefore, complete inhibition of both PD-L1 and PD-2 binding to PD-1 could be required to reverse ICB suppression.

**Figure 1.**
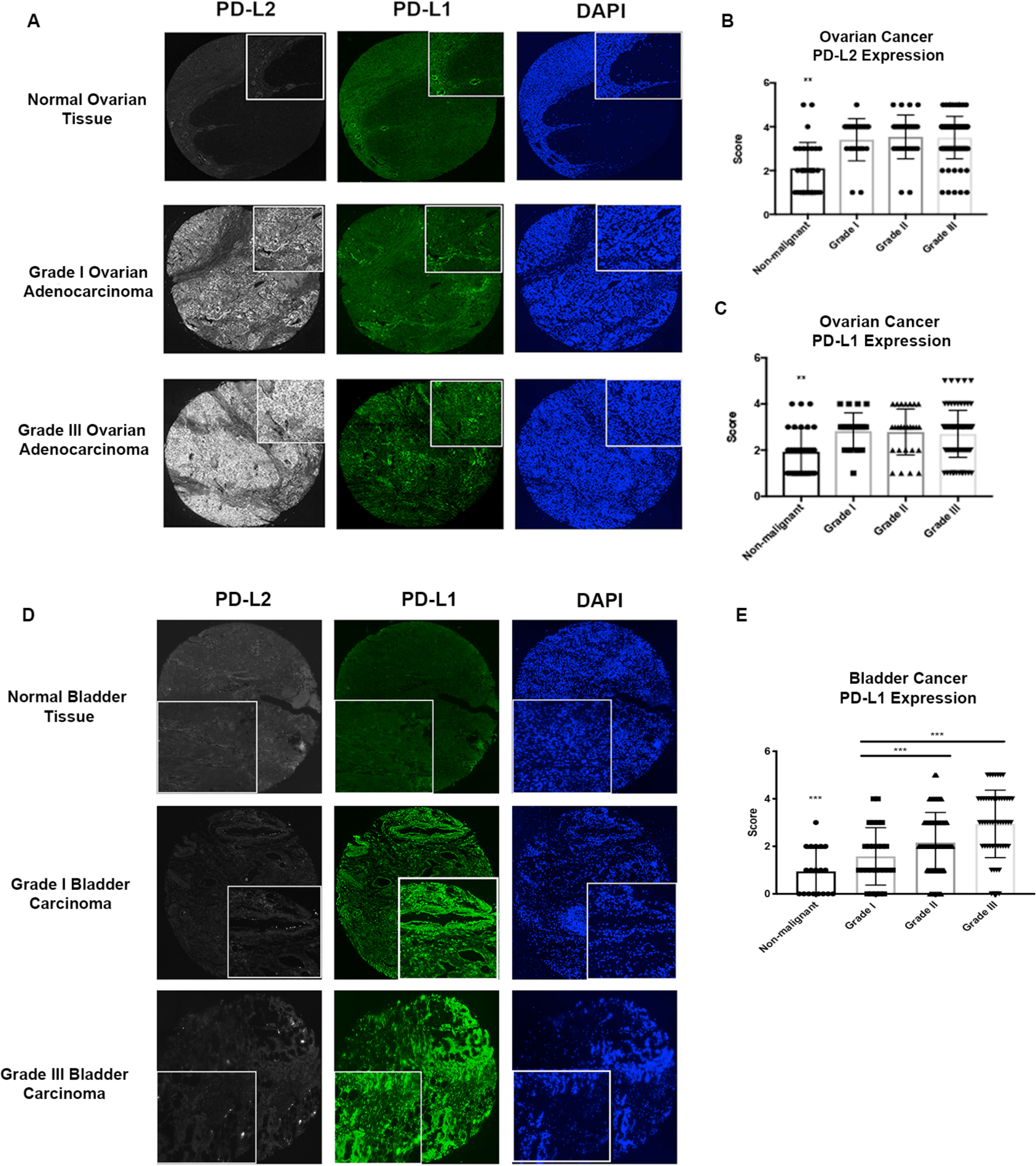
PD-L2, PD-L1 expression in ovarian and bladder cancer. (A) Representative images of ovarian cancer tissue microarray containing both normal and malignant samples (N=156) were stained with anti-human PD-L2 (gray), anti-human PD-L1 (green) and DAPI (blue) by fluorescent immunohistochemistry. The intensity of PD-L2 (B) and PD-L1 (C) were scored and stratified according to the tumor grade. (D) Bladder cancer tissue microarray was stained with anti-human PD-L2 (gray), anti-human PD-L1 (green) and DAPI (blue) by fluorescent immunohistochemistry with representative images shown (N=208). (E) PD-L1 expression in TMAs were scored and quantified according to tumor grade. Quantification data were plotted with mean and standard deviation calculated. One-way ANOVA were used for analysis comparing between tumor grades. P value *=<0.05, **=<0.01 and ***=<0.001.

### Engineering sPD-1 mutants with high binding affinity to PD-L1 and PD-L2

ICB is directed at inhibiting the binding of PD-1 on T cells or PD-L1 on tumor and stromal cells through the use of specific blocking antibodies. While these two groups of antibodies have been effective, PD-1 antibodies binding directly to T cells, potentially affecting its growth and viability and PD-L1 antibodies fail to inhibit the activity of PD-L2, which is also expressed on tumor and stromal cells. In addition, neither antibody has addressed the problem of the different affinities between PD-L1 and PD-L2 in binding PD-1. A different approach of antagonizing both PD-L1 and PD-L2 in a biological system is to utilize the soluble component of the PD-1 receptor (Fig. 2A).

**Figure 2.**
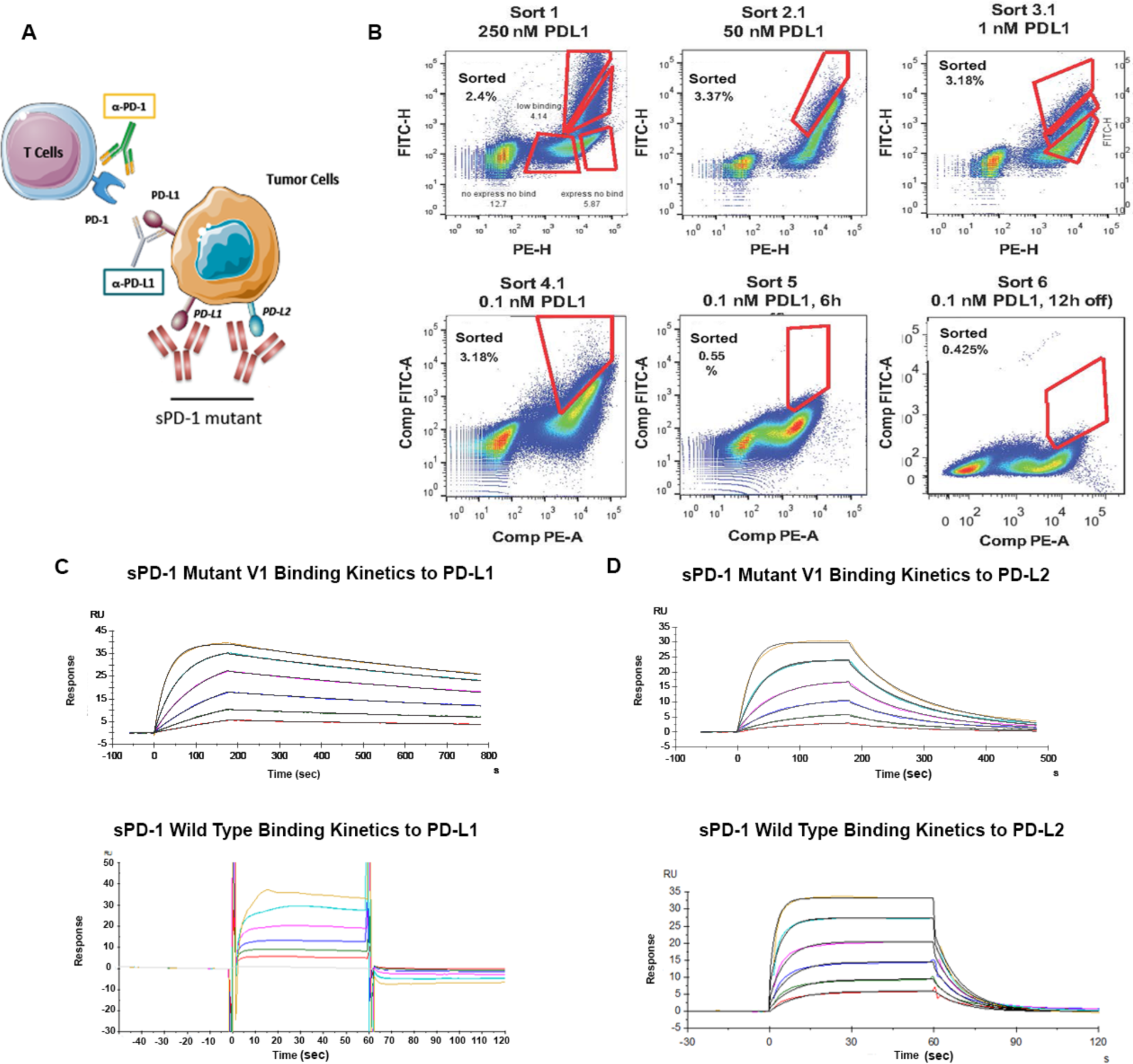
Engineering sPD-1 mutants with superior binding affinity to PD-L1 and PD-L2. (A) Conceptual illustration of sPD-1 mutant inhibiting the PD-1 signaling. (B) Representative flow cytometry dot plots showing clone selection pressure and gating strategies used for isolating high affinity PD-L1 binding clones. Clones with the strongest binding to PD-L1 were selected through sequential decrease of PD-L1 concentration (sort 1 to sort 3) followed by increase in incubation time (sort 4 and sort 6). (C) Binding kinetics of sPD-1 Mutant Version 1 (top) and wild-type PD-1 (bottom) binding to human PD-L1 was determined by BIAcore Surface Plasmon Resonance system. Each curve represents a single concentration of the analyte. (D) Binding kinetics of sPD-1 Mutant Version 1 (top) and wild-type PD-1 (bottom) binding to human PD-L2 was determined by BIAcore Surface Plasmon Resonance system. Each curve represents a single concentration of analyte

The native binding affinity of wild-type PD-1 to both of its ligands is in low micromolar range (28), making wild-type sPD-1 a reasonable antagonist for blocking PD-1/PD-L1 and to a lesser extent PD-1/PD-L2 binding compared with antibodies that are currently in the clinic. To develop a high affinity sPD-1 antagonist capable of neutralizing both PD-L1 and PD-L2, we generated a library of sPD-1 mutants by introducing random mutations using artificial nucleotides and error-prone PCR, throughout the complete extracellular domain of the PD-1 molecule (sPD-1). We used a yeast surface display based screening strategy that we have previously described to generate high affinity sAXL mutants (29, 30). The mutant library is displayed as fusion proteins on the yeast cell surface, and is screened by FACS to detect enhanced binding to fluorescently labeled PD-L1 ligands (Fig. 2B and Sup. Fig. 2). For FACS based selection, equilibrium binding sorts (sort rounds 1-3) were performed in which yeast clones were incubated with decreasing concentrations of hPD-L1 from 250 nM to 1 nM. These sort rounds were then followed by kinetic off-rate selection for sPD-1 mutants in sort rounds 4-6, where yeast clones were incubated with a fixed concentration of 0.1 nM PD-L1, after which the unbound PD-L1 was removed, and a ∼50 fold molar excess of soluble wild-type PD-1 was added to capture any remaining PD-L1 dissociated from the sPD-1 mutant binding complex. The length of the incubation time was 4 hrs (sort 4); 6 hrs (sort 5) and 24 hrs (sort 6). After six rounds of sorting, clones with high binding affinity were sequenced and further characterized (Table S1). Sequence analysis of frequently recurring mutations from the different sorts indicated that mutation of eight residues in the sPD-1 domain corresponding to Ser62, Ser87, Pro89, Asn116, Gly124, Ser127, Ala143 and Ala140 increased the binding affinity for PD-L1 and for PD-L2 (Table 2). Due to the fact that the mutation of Asn116 on sPD-1 corresponded to a N-glycosylation site, we generated a sPD-1 mutant with Asn116 unmutated (V2) and compared its biological features with the fully mutated version (V1). The sPD-1 mutants were fused to the Fc domain of human IgG4 with S228P mutation to minimize immunogenicity, improve plasma half-life, reduce the rate of renal clearance, and increase protein stability and solubility (31). The apparent affinities of both sPD-1 mutant V1 and V2 to human PD-L1 and PD-L2 were quantitatively measured using BIAcore® surface plasmon resonance technology. Compared to wild type sPD-1, an approximately 10,000-fold and 200-fold improvement in binding affinity towards hPD-L1 and hPD-L2 was observed, respectively (Table 1). Kinetics analysis of sPD-1 mutant V1 binding to PD-L1 and PD-L2 (Fig. 2C and 2D upper panel) at multiple concentrations (represented by each color-coded curve) demonstrated that the mutations in sPD-1 resulted in slower dissociation time upon binding to both ligands (Fig. 2C and 2D lower panel), a contributing factor towards enhanced binding affinity.

**Table 1:**
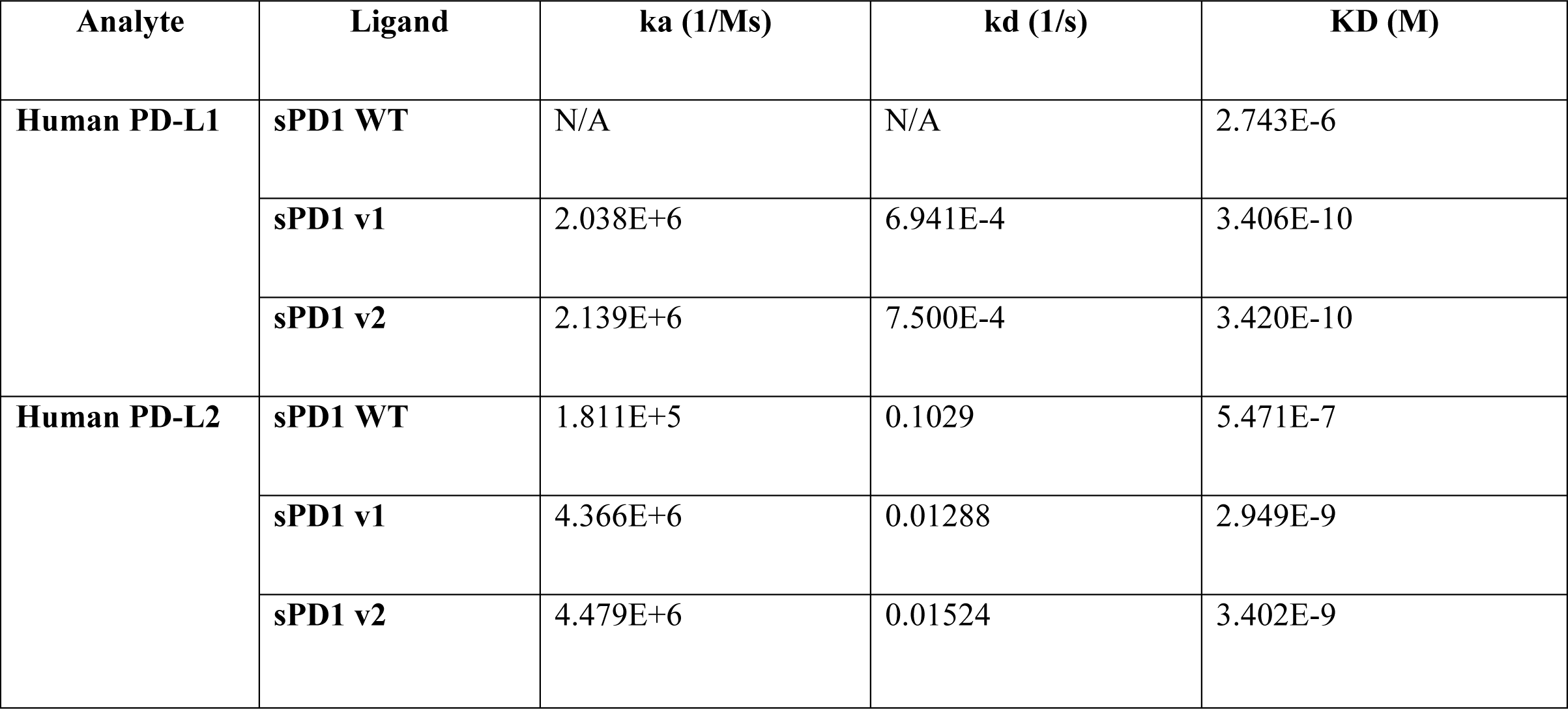
The apparent binding affinity between sPD-1 WT Fc, sPD-1 V1 Fc and sPD-1 V2 Fc in co-complex with human PD-L1 and PD-L2.

**Table 2:** Variations within the binding affinity and protein stability (in kcal/mol) for each mutation in co-complex with hPD-L1. Orange represents mutations outside of the binding interface; yellow represents of mutations within the binding affinity and white represent the N-glycosylation site.

| Residue | Original | Mutated | d Affinity | d Stability (solvated) |
| --- | --- | --- | --- | --- |
| 62 | SER | GLY | -1.37 | 0.5 |
| 87 | SER | GLY | 0.94 | 5.56 |
| 89 | PRO | LEU | 0 | -7.06 |
| 116 | ASN | SER | -0.02 | 0.82 |
| 124 | GLY | SER | -4.27 | -10.24 |
| 127 | SER | VAL | -0.12 | -1.82 |
| 132 | ALA | ILE | -10.94 | -11.9 |
| 140 | ALA | VAL | -0.77 | -25.89 |

### Computational based modeling to identify the structural basis for high-affinity binding

To determine the changes in structural conformation that result in an affinity increase between mutant hPD-1/hPD-L1 and mutant hPD-1/hPD-L2 compared to wild-type, we performed computational model simulations. Wild-type human PD-1/PD-L1 complex structures obtained from Protein Data Bank (PDB No. 4ZQK and 5IUS) were used as backbones for mapping amino acid mutations identified in the sPD-1 mutants. Sequence alignment of both hPD-1 structure No. 4ZQK and No. 5IUS is shown in Sup. Fig 4A. The sPD-1 mutant structure was modeled using the Residue Scanning module in Schrodinger Suite. The crystal structure of sPD-1 and hPD-L1 are shown in blue and green respectively with the 8 mutations (Ser62, Ser87, Pro89, Asn116, Gly124, Ser127, Ala132 and Ala140) shown in orange sticks (Fig. 3A). Based on structural modeling data, we identified residues G124S, S127V and A132I as mutations within the binding interface, and S62G, S87G, P89L, and A140V outside of the binding interface. Mutations G124S and A132I each make one additional hydrogen bond with Tyr 123 and Gln 66 of PD-L1 respectively (Fig. 3B left and right panel). Mutation A132I on sPD-1 resulted in an increase in the protein surface complementarity with hPD-L1 from 0.72 to 0.85 (Fig. 3C, Sup. Fig. 4B). To further investigate how these mutations influence the overall binding between sPD-1 mutations and hPD-L1, the variation of the binding affinity and complex stability before and after individual mutation were calculated and the results are listed in Table 2. We also calculated the variation in binding affinity and complex stability for grouped mutations within and outside of the binding interface (Table S2). In both Table 2 and S2, a higher negative value is indicative of increased binding affinity and stability. Overall, the interface mutations contribute to both binding affinity and protein stability as expected, and the mutations outside of the binding interface lead to increased stability.

**Figure 3.**
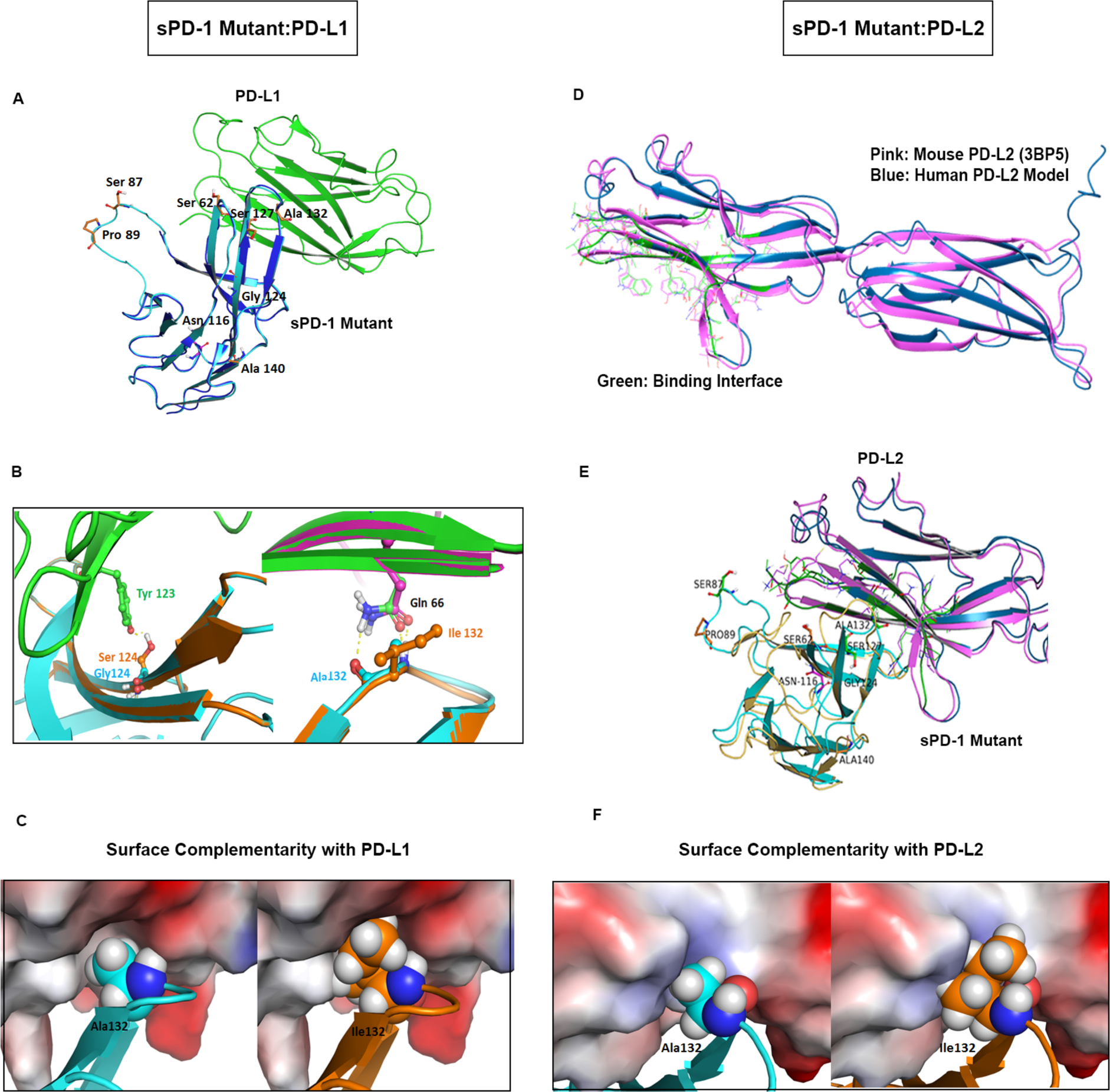
Computational modeling based structural analysis of sPD-1 Mutants in co-complex with human PD-L1 and PD-L2. (A) Overlay of human PD-1 model in complex with human PD-L1 (PDBID:4ZQK) are shown the blue and green respectively. The model of PD-1 with missing loop fixed is shown in cyan and the 7 residues mutated on sPD-1 are shown in orange sticks, the N-glycosylation site (Asn116) is shown in magenta sticks. (B) Left panel demonstrates the sPD-1 mutation G124S (orange) makes a hydrogen bond with PD-L1 Tyr123 (green and red sticks). Wild-type PD-1 structure is shown in cyan. Right panel shows mutation A132I makes one more hydrogen bond with PD-L1 Gln166. Wild-type PD-1 structure (cyan), mutated PD-1 structure (orange). Crystal PD-L1 structure is shown in pink and PD-L1 co-complex with sPD-1 mutant is shown in green. (C) Comparison of surface complementarity of mutation A132I. PD-L1 binding site is marked to reveal electrostatic potential surface. Red indicates negative electrostatic potential, blue indicates positive electrostatic potential and grey indicates hydrophobic regions. Wild-type PD-1 is shown in cyan cartoon and balls (left panel); mutated PD-1 is shown in orange cartoon and balls (right panel). (D) Alignment of proposed human PD-L2 with mouse PD-L2 (PDBID: 3BP5). The interface residues are marked green within the dark blue human PD-L2 model and the crystal structure of mouse PD-L2 labeled pink. (E) The overlay of proposed Human PD-1 with mutations (cyan) in complex with human PD-L2 (dark blue) and mouse PD-1 (yellow) in complex with mouse PD-L2 (pink). (F) Comparison of surface complementarity of mutation A132I with PD-L2 with same color annotation as (C).

**Figure 4.**
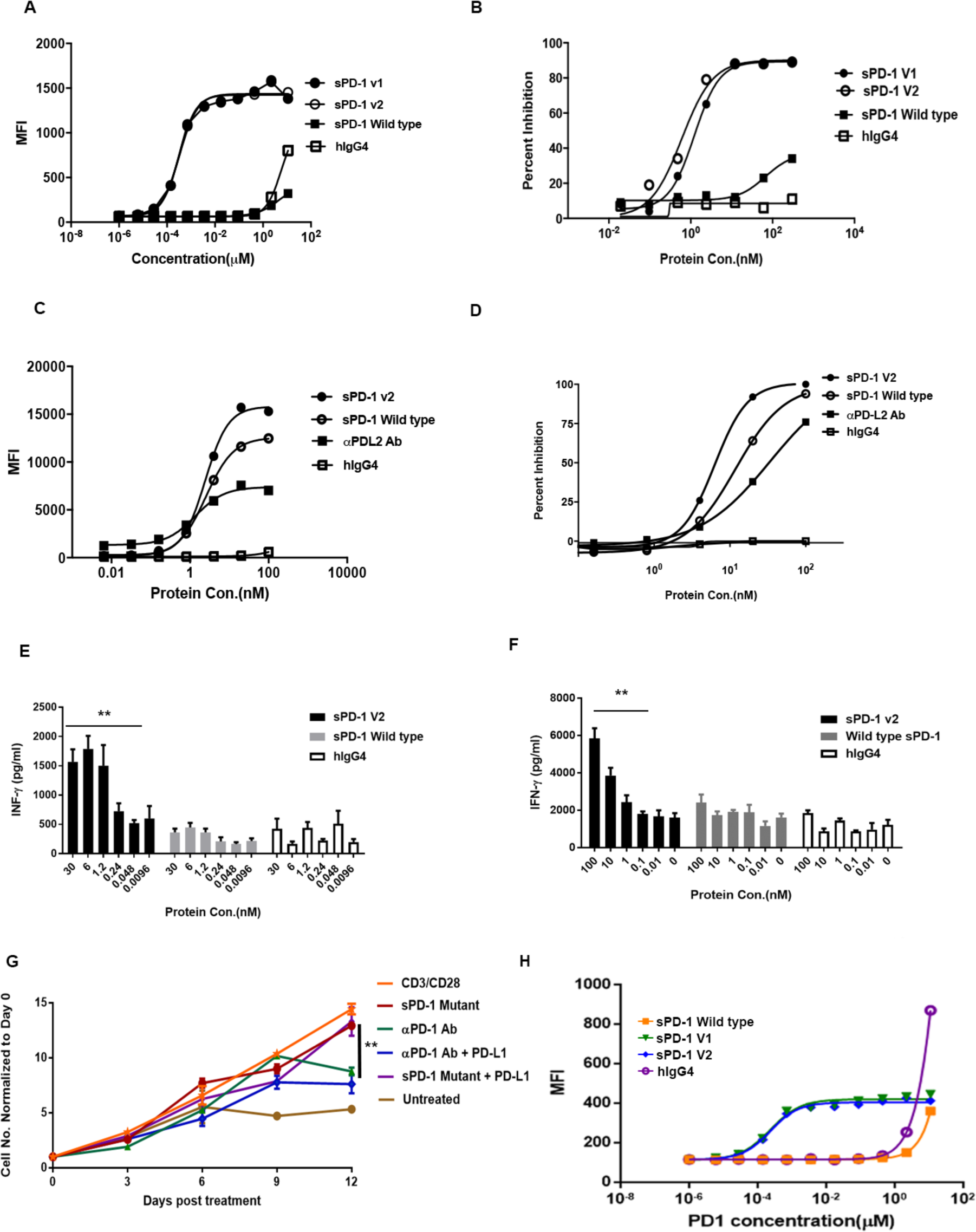
sPD-1 mutants demonstrated superior capability in blocking PD-L1 and PD-L2 mediated activities in a ligand-dependent manner without affecting T cell viability. (A) Binding of wild-type PD-1, hIgG4 and sPD-1 mutants V1 and V2 to MC38 cells with human PD-L1 knock-in (MC38-hPD-L1). Figures A-D are all presented as one of the two independent experiments shown here with individual points representing the mean of two technical repeats. (B) Cell-based receptor-blocking assay showing inhibition of Hep3B-hPD-L1 binding to biotin conjugated PD-1 wild-type in competition with sPD-1 wild-type, hIgG4 and sPD-1 mutants. (C) Binding of wild-type PD-1, hIgG4 and sPD-1 mutant V2 to MC38-hPD-L2. (D) Cell based receptor-blocking assay showing inhibition of Hep3B-hPD-L2 binding to biotin conjugated PD-1 wild-type in competition with sPD-1 wild-type, hIgG4 and sPD-1 mutants. (E) T cell activation in the presence of Hep3B-hPD-L1 cells when incubated with sPD-1 mutant, sPD-1 wild-type and hIgG4. IFN-γ levels measured as a marker of T cell activation. Error bars represent the mean and standard deviation of technical triplicate. Experiment was conducted twice with PBMCs isolated from different donors. (F) T cell activation in the presence of Hep3B-hPD-L2 cells upon incubation with sPD-1 mutant, sPD-1 wild-type and hIgG4. T cell activity is measured by IFN-γ . Error bars represent the mean and standard deviation of technical triplicate. Experiment was conducted twice independently with PBMC isolated from different donors. (G) T cell proliferation over-time in the presence of sPD-1 mutant V2 and αPD-1 antibody with or without hPD-L1 added.(H) Binding kinetics between sPD-1 mutants and mouse PD-1 in MC38 parental cells. Each data point represents the mean and standard deviation of technical duplicate. Experiment was repeated with T cells isolated from a different donor. Statistical analysis was conducted with One-way ANOVA for comparing between treatment groups and repeated ANOVA for changes over time. P value *=<0.05, **=<0.01.

Given that the hPD-L2 protein structure was not resolved until very recently (32), a model of the hPD-L2 was simulated based on available mPD-L2 structures (Sup. Fig. 5A). The sequence identity between hPD-L2 and mPD-L2 is about 72% (Sup. Fig. 5A). The overlay of the proposed hPD-1/PD-L2 model and murine PD-1/PD-L2 crystal structure (PDBID:3BP5) is shown in Fig. 3D and Sup. Fig.5B. The proposed co-complex structure between sPD-1 mutations with the proposed human PD-L2 is shown in Fig 3E. Compared to the proposed wild-type hPD-1/hPD-L2 co-complex, mutation A132I created an additional hydrogen bond with PD-L2, resulting in increased surface complementarity (Fig. 3F and Sup. Fig 6A). The calculation of the binding affinity variation and protein stability was also performed for single mutations and grouped mutations within and outside of the binding interface (Table 3 and Sup. Fig. 6B). Importantly, the A140V mutation, in sPD-1, which lies outside of the binding interface, is a significant contributor towards enhanced binding affinity by increasing the overall stability of the sPD-1/PD-L1, PD-L2 co-complexes. This observation further emphasizes the importance of a non-biased screening approach for identifying mutations outside of a binding interface for improving the overall stability of the protein-protein interaction.

**Figure 5.**
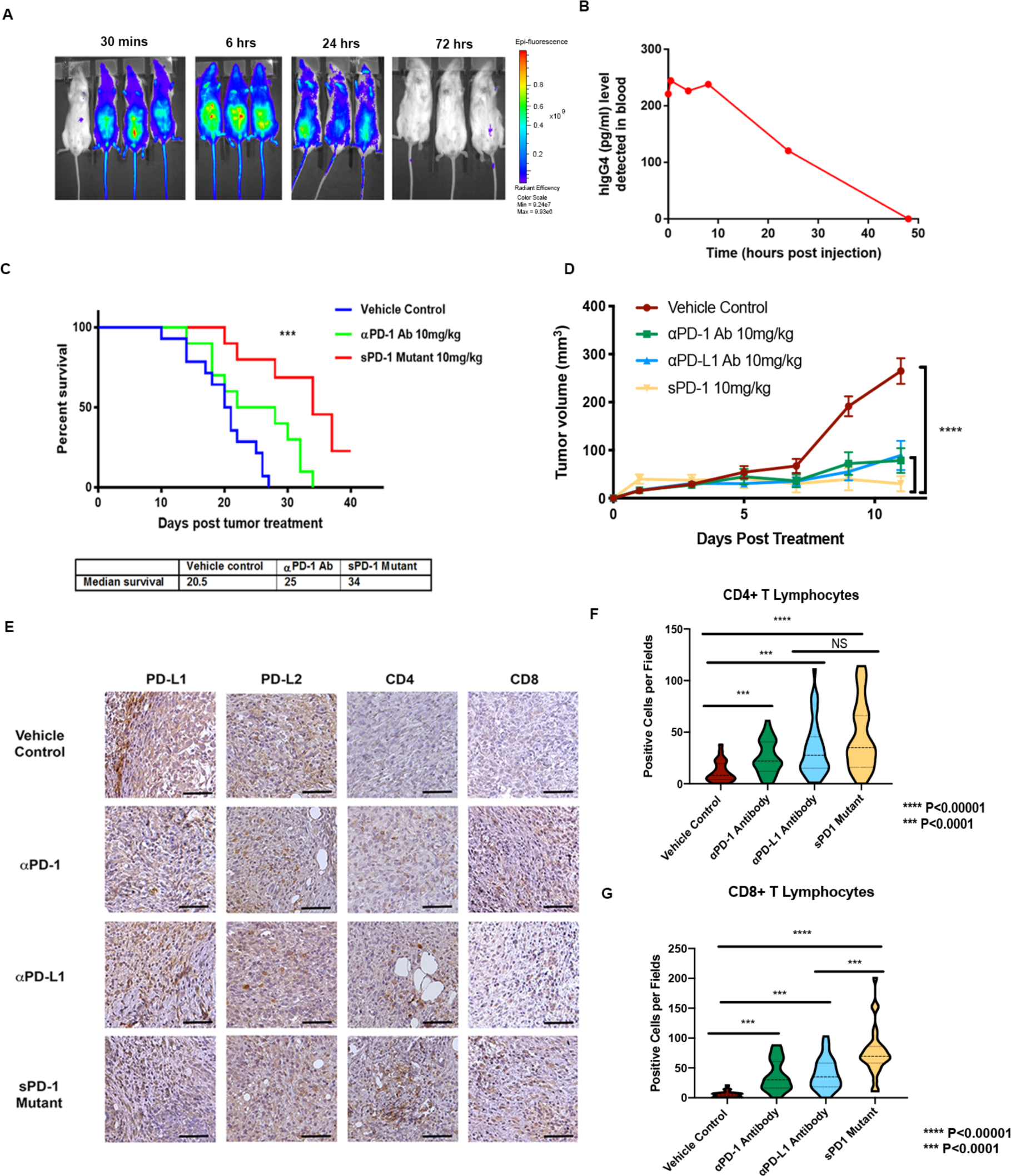
sPD-1 mutant inhibits tumor growth in mouse tumor models of ovarian cancer. (A) Bio-distribution of sPD-1 mutant labeled with Alexa Fluor® 488 after a single dose of 10mg/kg were imaged over-time. (B) Bio-distribution of sPD-1 mutant in mouse serum after a single dose of molecule at 10mg/kg detected with ELISA against human IgG4. Each data points represent the mean of two animals collected at the same time point. (C) Kaplan Meier survival plot of C57B/6 mice orthotopically inoculated with ID8 mouse ovarian tumor cells treated with vehicle control, anti-mouse αPD-1 blocking antibody 10mg/kg and sPD-1 mutant 10mg/kg. Animals terminated upon the development of ascites. Median survival of each experimental group listed below. (D) Subcutaneous UPK10 ovarian tumor growth over-time in C57B/6 mice treated with vehicle control, sPD-1 Mutant, αPD-1 antibody and αPD-L1 antibody. (E) The expression of PD-L1, PD-L2, CD4 and CD8 positive cells in UPK10 tumors post-treatment was analyzed by IHC staining. Scale bar 25μm. (F) Violin plot of CD4+ T lymphocyte infiltration into UPK10 ovarian tumors post treatment. (G) Violin quantitative plot of CD8+ T lymphocyte infiltration into UPK10 ovarian tumors post treatment. Statistical analysis was conducted using One-way ANOVA for comparing between treatment groups and repeated ANOVA for changes occur over-time. Kaplan Meier estimator was calculated for survival curves. P value *=<0.05, **=<0.01. ***=<0.001.

**Figure 6:**
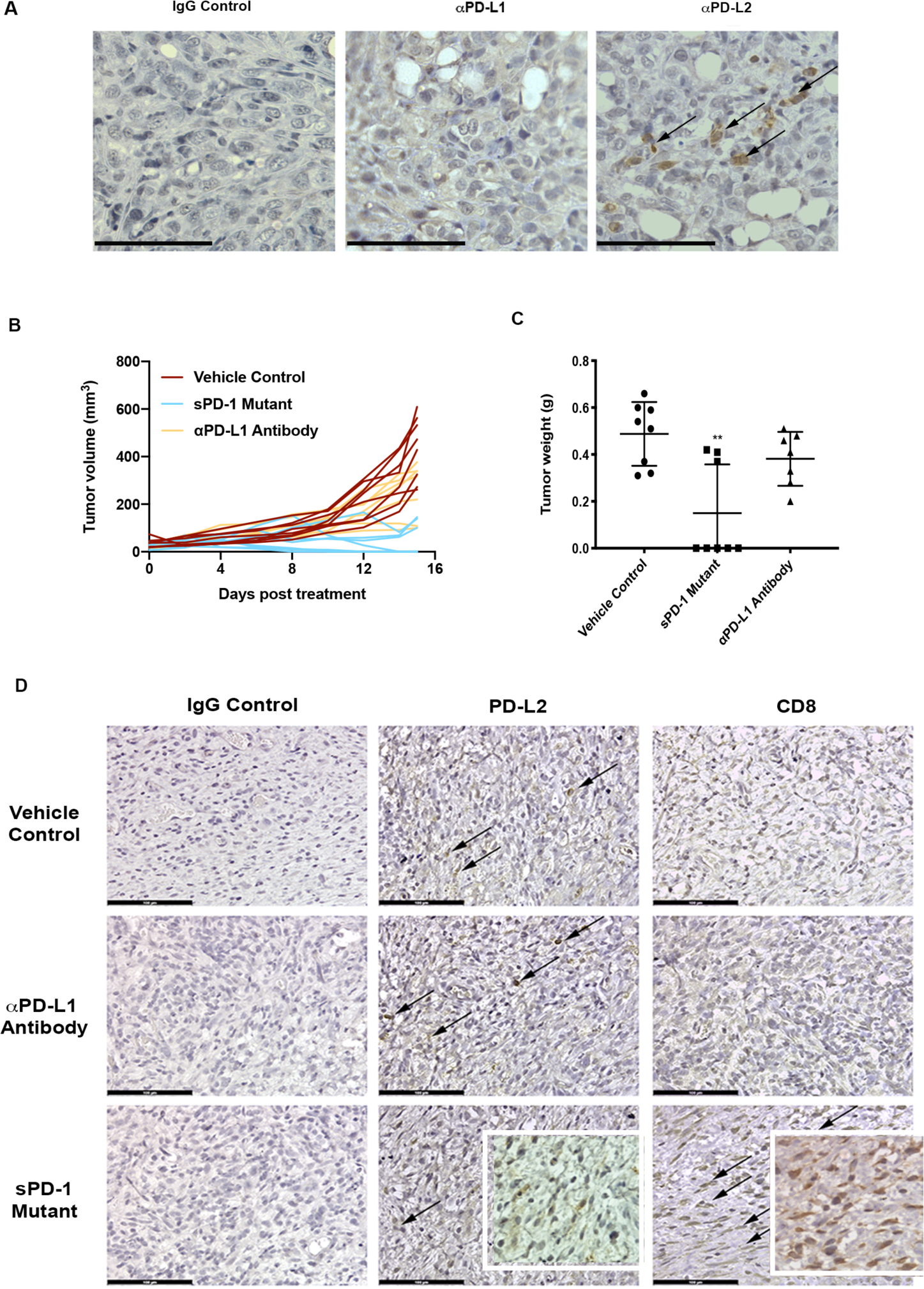
sPD-1 mutant treatment suppresses tumor growth in a ovarian cancer model that is PD-L2 dependent. (A) ID8 PD-L1 CRSIPR tumors inoculated into PD-L1 KO mice. Tumor stained for PD-L1 and PD-L2 expression. Scale bar 50μm. (B) Subcutaneous ID8 PD-L1 CRSIPR tumors growth in C57B/6 PD-L1 KO mice. Mice were treated with vehicle control, anti-mouse αPD-L1 blocking antibody and sPD-1 mutant V2. (C) Total tumor weight of ID8 PD-L1 CRISPR KO tumors at the time of termination. (D) IHC staining of PD-L1 and PD-L2 in ID8 PD-L1 CRISPR KO tumors treated with vehicle control, anti-mouse αPD-L1 blocking antibody 10mg/kg and sPD-1 mutant 10mg/kg. Scale bar 100μm. Statistical analysis conducted using One-way ANOVA for comparing between treatment groups and repeated ANOVA for changes over time. P value *=<0.05, **=<0.01. ***=<0.001.

**Table 3:** Variations within the binding affinity and protein stability (in kcal/mol) for each mutation in co-complex with hPD-L2. Orange represents mutations outside of the binding interface; yellow represents of mutations within the binding affinity and white represent the N-glycosylation site.

| Residue | Original | Mutated | d Affinity | d Stability (solvated) |
| --- | --- | --- | --- | --- |
| 62 | SER | GLY | 0.3 | 4.21 |
| 87 | SER | GLY | 0 | -0.68 |
| 89 | PRO | LEU | 0.28 | -4.71 |
| 116 | ASN | SER | 0 | 0.85 |
| 124 | GLY | SER | -3.06 | 6.47 |
| 127 | SER | VAL | 1.65 | -4.42 |
| 132 | ALA | ILE | -9.47 | -0.55 |
| 140 | ALA | VAL | 0.25 | -11.76 |

### sPD**-1** mutants disrupts PD-L1 and PD-L2 binding to PD-1, result in T cell activation

The sPD-1 mutants (V1, V2) and wild-type sPD-1 were transiently expressed using HEK293 cells. Purified protein and endotoxin levels were tested by SDS-PAGE, LC/MS and endotoxin assays (Sup. Fig. 7A-F). To demonstrate that the sPD-1 mutants can disrupt the interactions between PD-L1 or PD-L2 and PD-1 at the cellular level, we assessed the binding activity of sPD-1 mutants to PD-L1 using cell lines with a knock-in human PD-L1. We performed our studies in MC38 hPD-L1 knock-in cells, because our attempts to generate a hPD-L1 knock-in using murine ovarian cancer cell ID8 and UPK10 were not successful. In this model, FACs based binding analysis showed similar level of improvement in the binding affinities of sPD-1 V1 and V2 mutants towards hPD-L1 expressed on MC38 cells, indicating that the mutation at the Asn116 N-glycosylation site is not necessary for the enhancement of binding affinity. Consistent with previously reported literature, wild-type sPD-1 binds to hPD-L1 with low affinity and no fluorescent signal was detected up to µM concentration (Fig. 4A) To further examine the ability of sPD-1 mutants to block binding between endogenously expressed hPD-L1 and PD-1, we performed Receptor Binding Assays (RBA) using sPD-1 mutants, wild-type sPD-1, and hIgG4 controls. Both sPD-1 mutants showed superior inhibition of PD-L1 mediated receptor binding towards PD-1 when compared to wild-type sPD-1 and hIgG4 (Fig. 4B). To demonstrate that the sPD-1 mutant is capable of binding to PD-L2 and block PD-L2 mediated PD-1 activation, we overexpressed human PD-L2 in mouse MC38 cells (MC38-hPD-L2) and human hepatocellular carcinoma Hep3B cells (Hep3B-hPD-L2) to generate stable cell lines (Sup. Fig. 9 and 10). Given that no differences in binding were detected between sPD-1 V1 and sPD-1 V2, we decided to continue our study with sPD-1 mutant V2 without the Asn116 mutation. The sPD-1 V2 mutant exhibits the strongest binding towards endogenous hPD-L2 cells while both wild-type sPD-1 and the commercially available, generic αPD-L2 antibody showed a 25% and 50% reduction in mean fluorescent intensity (MFI) at the same concentrations (Fig. 4C). The sPD-1 mutant also exhibits enhanced ability to block PD-L2 mediated activation of PD-1 in receptor binding assays (Fig. 4D). The binding analysis was also repeated in Hep3B-hPD-L2 with similar results (Sup. Fig. 8A). We then investigated whether the sPD-1 mutant can functionally inhibit PD-1 signaling in activated T cells. Human PBMCs were collected from healthy donors and incubated with Hep3B-OS8-hPDL1 or Hep3B-OS8-hPDL2 cells. The sPD-1 mutant, wild-type PD-1, and IgG4 were then added in increasing concentrations and T cell activity was measured by Interferon-γ secretion. We found that the sPD-1 mutant activated T-cells in the presence of endogenous PD-L1 or PD-L2 in a dose-dependent manner (Fig. 4 E and F), supporting the concept that the sPD-1 V2 can functionally stimulate T-cell activity through neutralizing the interaction between PD-L1/PD-L2 and PD-1. Since prolonged stimulation of T-cell activity through direct binding using αPD-1 antibodies and PD-1 receptors on T-cells can potentially affect T-cell functionality and viability over time, our sPD-1 mutant was designed to primarily target cancer cells expressing high levels of PD-L1 and PD-L2, thus aiming to preserve T-cell longevity and functionality. To test this hypothesis, activated human T-cells (through stimulation with CD3/CD28 cocktail) were treated with either αPD-1 antibody or sPD-1V2 mutant in the presence or absence of PD-L1 over a period of 12 days. Continuous treatment with αPD-1 slowed T-cell proliferation over-time while sPD-1 mutant treatment did not affect T-cell growth, showing a similar growth rate as the CD3/CD28 stimulated T-cells. In contrast, naive T cells did not proliferate in the absence of CD3/CD28 stimulation (Fig. 4G). These observations indicate that our sPD-1 mutant is capable of binding and blocking the PD-1/PD-L1 and PD-L2 signaling cascade and stimulating T-cell activity while preserving T-cell viability and functionality. Since human PD-1 is known to interact with mouse PD-L1, and the PD-1 molecule is conserved between murine and human genomes (33), we validated the binding of sPD-1 mutants to mouse PD-L1 in wild-type parental MC38 cells, which only express mouse PD-L1 and PD-L2. An approximate 4-fold increase in binding signal was detected with sPD-1 V1 and V2 compared to wild-type sPD-1 (Fig. 4H). This species cross reactivity allows us to test the sPD-1 mutants in a syngeneic manner in murine models of human malignancies.

### sPD-1 mutant demonstrates superior anti-tumor efficacy in syngeneic ovarian and other mouse cancer models

The therapeutic potency of the sPD-1 mutant was evaluated in multiple syngeneic mouse tumor models of both ovarian and other sites. Mutant sPD-1V2 which lacks the N-glycosylation site (Asn116) mutation was used for all *in vivo* studies. To evaluate the pharmacokinetics of the sPD-1 mutant *in vivo,* a single dose of fluorescently-labeled sPD-1 mutant at a concentration of 10mg/kg was injected into mice and imaged at 30 minutes, 6 hours, 24 hours and 72 hours post-treatment (Fig. 5A). We also used an ELISA assay to detect the human IgG4 component of the sPD-1 mutant molecule as a direct means to monitor drug levels in the serum over time (Fig. 5B). Both fluorescent images and ELISA analysis indicated that the half-life of the sPD-1 mutant molecule is approximately 24 hours, providing the rationale for *in vivo* dosing at 10 mg/kg every 48 hours.

To determine the efficacy of sPD-1 compared to an-anti-PD-1 antibody, we intraperitoneally inoculated C57/Bl6 mice with ID8 mouse ovarian cancer cells and treated with vehicle control, anti-mouse αPD-1 antibody or sPD-1 mutant every 48 hours. Animals were monitored for survival and terminated upon developing ascites. A significant improvement was found in the survival of mice treated with the sPD-1V2 compared to the vehicle control group (34 days vs. 20.5). The αPD-1 treated group showed an intermediate response between the control and sPD-1V2 treated groups (Fig. 5C). A subset of mice containing ID8 tumors from each of the treatment group was sacrificed on day 20 post-treatment and analyzed for PD-L1, PD-L2, CD4 and CD8 expression immunohistochemically (Sup Fig 11A). Tumors in all groups stained positive for PD-L1 and PD-L2 expression, but significantly higher numbers of CD4+ and CD8+ T cells were observed infiltrating the tumors treated with sPD-V2 compared to vehicle control and αPD-1 treated group (Sup. Fig. 11B, C). A second syngeneic mouse ovarian cancer model UPK10 was tested. Animals treated with sPD-1 mutant, αPD-1 or αPD-L1 antibodies. A significant reduction in tumor growth was observed in all three treated groups, but was most pronounced in sPD-1 mutant treated animals (Fig. 5D). IHC Stained UPK10 tumors showed a positive correlation between antitumor activity of the sPD-1 mutant and increased CD4+ and CD8+ T cell infiltration. While TILs were also observed in both αPD-1 and αPD-L1 treated groups, they were further elevated in the sPD-1 mutant treated tumors (Fig. 5E-G). No significant changes in animal body weight were observed throughout the experiment (Sup. Fig. 11D). The therapeutic potential of the sPD-1 mutant was also studied in non-ovarian tumor models. MC38-hPDL1 colorectal cancer cells were inoculated subcutaneously in C57/B6 mice and treated with vehicle control, sPD-1 mutant or αPD-L1 antibody (atezolizumab). A significant reduction in tumor growth was observed in both sPD-1 mutant and αPD-L1 (atezolizumab) treated tumors (Sup. Fig. 12A, C). Compared to αPD-L1 treatment, tumors treated with the sPD-1 mutant showed superior efficacy in suppressing tumor growth and prolonged survival with no severe toxicity defined by change in body weight (Sup. Fig. 12B, D). The ability for the sPD-1 mutant to antagonize PD-L2 mediated tumor growth was further demonstrated in *in vivo* models inoculated with MC38 tumors overexpressing hPD-L2 and compared to tumors treated with αPD-1 antibody pembrolizumab. Significant delays in tumor growth were observed in tumor-bearing mice treated with sPD-1 mutant or pembrolizumab. Although statistically not significant, there was a trend towards better suppression of tumor growth in the sPD-1V2 treated group compared to the pembrolizumab treated group that may warrant further investigation (Sup. Fig. 13A). The antitumor activity of the sPD-1 mutant was further tested in mouse B16/OVA melanoma model. Melanoma tumors treated with sPD-1 mutants resulted in reduced tumor growth compared to vehicle treated groups (Sup. Fig. 13B). In separate studies, TILs were analyzed in mice bearing B16/OVA (Sup. Fig. 14A) and MC38 (Sup. Fig. 14B, C) tumors treated with sPD-1 mutant or αPD-1 antibody. While both treatments resulted in increased infiltration of CD4^+^, CD8^+^ cytotoxic T-cells, the most significant changes were observed in the sPD-1 mutant treatment groups. Tumor infiltrating NK cells and Macrophages were also analyzed but no significant differences were detected between treatment groups (Sup. Fig. 14D, E). Collectively, these studies demonstrate strong antitumor efficacy of the sPD-1 mutant in multiple syngeneic mouse tumor models, and such antitumor activity is partly driven by enhanced infiltration of TILs.

### Efficacy of sPD-1 mutant in a PD-L2 driven animal model of human ovarian cancer

To demonstrate the requirement of PD-L2 in facilitating PD-1 mediated tumor growth. We genetically deleted PD-L1 in ID8 ovarian tumor cells and MC38 colorectal cells using CRISPR-CAS9 system and seeded tumor cells in PD-L1 knockout mice, eliminating PD-L1 expression in both the tumor and the host (Fig. 6A and Sup. Fig. 15). Interestingly, MC38 PD-L1 KO model fail to form tumor in PD-L1 KO mice while ID8 PD-L1 KO model did with higher number of tumor cells seeded. In ID8 PD-L1 KO model, blocking PD-L2 interaction with PD-1 through administering sPD-1V2 led to significant tumor regression, and tumors treated with αPD-L1 antibody remained largely unresponsive (Fig. 6B, C). When ID8 tumors were stained for both PD-L2 and CD8 expression, all tumors regardless of treatment groups showed positive PD-L2 expression. However, only tumors treated with sPD-1V2 saw infiltration of CD8+ T cells into the tumor (Fig. 6D), demonstrating the importance of blocking PD-L2 mediated activation of PD-1.

In this study, we demonstrate the clinical relevance of high PD-L2 expression in human ovarian cancer, a possible contributing factor leading to poor response towards ICB. We developed a sPD-1V2 molecule which blocks the activity of PD-L1 and PD-L2 to PD-1 with ultra-high affinity. Structural and biological analysis of the sPD-1V2 strongly supports our hypothesis that improvement in the binding affinity enhances PD-1 inhibition by blocking both PD-L1 and PD-L2 ligands. Thus, our enhanced sPD-1V2 represents a new therapeutic approach for treating ovarian cancer, which express both PD-L1 and PD-L2.

## Discussion

To date, the single agent activities of anti-PD-1 and anti-PD-L1 inhibitors in some cancers have being disappointing (24, 26). The recent JAVELIN Ovarian 100 trial, which evaluated a αPD-1 antibody (avelumab) in combination with and/or following platinum-based chemotherapy in 998 previously untreated patients, found a poor overall response. Similar outcomes have been also reported in gastric, and brain malignancies (24, 25, 34, 35). The immunosuppressive TME in these cancer types is particularly challenging to overcome. In particular, tumor cells and surrounding stromal tissue in the TME express both PD-L1 and PD-L2 to facilitate tumor escape, preventing both the recruitment and activation of TILs (36, 37). Therefore complete inhibition of the PD-1 signaling pathway through neutralizing both of its ligands is necessary to boost T cell mediated killing. Currently in the clinic, inhibition of the PD-1 signaling is achieved through blocking the PD-1 receptor or PD-L1. PD-L2 still remains therapeutically unexplored in the context of ICB with only one clinical candidate, AMP-224, which comprises a Fc fusion with the extracellular domain of PD-L2 to act as ligand trap for PD-1 receptor (38), The lack of therapeutic interest in targeting PD-L2 is predominantly based on the belief that the tissue expression of PD-L2 is low and irrelevant during cancer progression and does not consider the possibility that PD-L2 expression is most likely dependent on TME stimulation. It is also widely believed that blocking PD-1 receptor therapeutically is sufficient to inhibit the interaction between PD-1 receptor and both of its ligands. However, emerging evidence from our laboratory and others have found strong clinical and biological evidence that in some cancer types, PD-L2 expression is up-regulated by tumors and its TME to promote survival and immune evasion (39–41). In addition, PD-L2 binds to PD-1 receptor at 6- to10- fold higher binding affinity compared to PD-L1, which represents significant competition with therapeutic anti-PD-1 antibodies for receptor binding and engagement, leading to treatment escape.

Unlike PD-L1, which is expressed in abundance on a variety of both malignant and normal cells, it is thought that basal PD-L2 expression is low on tumor cells and restricted to very few immune cell types of the myeloid lineage. Dendritic cells (DCs) and macrophages produce Th2 cytokines such as IL-4 and drive PD-L2 expression, leading to T cell suppression and immune escape (40). However, it has been reported that T cells can also up-regulate PD-L2 through IL-4 mediated signaling, which can also result in inhibition of T cell activity (40, 42). In the context of cancer, studies have reported the induction of PD-L2 expression in both cancer and tumor-associated stromal components upon malignant transformation, which could also serve as a mechanism for immune evasion (41). Our data suggests that high PD-L2 expression is often associated with cancers with low clinical response towards anti-PD-1 and anti-PD-L1 therapies. In particular, ovarian cancer showed elevated PD-L2 in cancer tissue and its surround stroma, but is not detectable in normal ovary tissue, supporting the concept that the expression of PD-L2 is inducible by microenviromental stimulation during tumor progression, most likely as a counter-measure towards immune surveillance. Therefore, adequate inhibition of both PD-L1 and PD-L2 in cancer is necessary for maximal blockade of the PD-1 signaling pathway.

The reported increase in the apparent binding affinity between PD-L2 and PD-1 compared to PD-L1, suggests that the presence of abundant PD-L2 can undermine the therapeutic activity of αPD-1 antibodies through competing for PD-1 receptor binding. Our strategy to overcome this affinity barrier is to develop sPD-1 mutants with superior binding to both PD-L1 and PD-L2. Using a yeast surface display based platform and unbiased mutation based directed evolution approaches, our sPD-1 mutant clones displayed significant affinity enhancement towards both PD-L1 and PD-L2. Unlike directed mutagenesis approaches that only mutate residues at the receptor-ligand binding interface, our unbiased mutation strategy identified three mutations within the binding interface of sPD-1, as well as four mutations outside of the sPD-1 binding domain. In particular, the mutation on Ala140 to Val located outside of the binding interface resulted in improved overall stability to the PD-1/PD-L1 and PD-1/PD-L2 co-complexes, providing structural evidence for the importance of searching beyond the binding interface for affinity enhancement. Interestingly, while there were three studies previously published that also generated high-affinity binding sPD-1 clones, none have reported mutations at A140, nor analyzed their mutant molecules for binding towards PD-L2 (32, 43, 44). Compared to the binding affinity of commercially available anti-PD-L1 antibody atezolizumab (TENCENTRIQ), our sPD-1 mutants have enhanced binding affinity to PD-L1 (kd= 3.406E^-10^ vs. kd = 9.96E^-09^ M) under similar experimental conditions (28). In addition, our sPD-1 mutants have enhanced binding towards PD-L2, whereas atezolizumab does not bind to PD-L2 (45). Therefore, sPD-1V2 is well characterized in regards to its dual-ligand neutralizing capabilities.

We tested the sPD-1 mutant in two syngeneic ovarian cancer models as well as other cancer models to address two questions: 1) Can the sPD-1 mutant outperform both αPD-L1 and αPD-1 antibodies to control tumor growth based on its enhanced ability to neutralize both PD-1 ligands? 2) Can sPD-1 mutant affect tumor growth in ovarian cancer, which is solely driven by PD-L2? We found that our sPD-1 mutant out-performed both αPD-L1 and αPD-1 antibodies in syngeneic models of ovarian and other cancer types showing improved CD4^+^ and CD8^+^ TIL infiltration. More importantly, when PD-L1 was genetically ablated from both the cancer cells and inoculated into PD-L1 knockout mice, only sPD-1 mutant but not αPD-L1 antibody treated animals exhibited antitumor activity. However, it is noteworthy that while we tested both MC38 PD-L1 CRISPR KO model and ID8 CRISPR KO model, MC38 PD-L1 KO model failed to form tumor, that is consistent with previous report that MC38 tumor growth largely relies on host PD-L1 expression (46). Therefore the anti-tumor activity we observed in MC38 *in vivo* study is likely due to the enhanced PD-L1 binding and signaling neutralization contributed through sPD-1 mutants. For ID8 CRISPR KO model, higher tumor cell number were needed for tumor establishment and delayed tumor growth was also observed compare to wild-type ID8 mouse model, indicating that for this model, both PD-L1 and PD-L2 expression enhances tumor growth and immune evasion, supporting our approach to develop a dual PD-L1 and PD-L2 blockade.

In this study, we identified PD-L2 as a factor that is associated with poor clinical response towards PD-1 inhibitors in ovarian cancer. By screening for amino acid mutations both within and outside of the sPD-1 binding interface, we developed sPD-1 mutants with superior binding affinities to trap both PD-L2 and PD-L1. Given that improving the response rate of ICB is an urgent unmet need for ovarian cancer patients, data presented in this study provided justification for using high-affinity sPD-1 mutant as an alternative to PD-1 or PD-L1 therapeutic antibodies to achieve superior therapeutic efficacy in cancers expressing both PD-L2 and PD-L1.

## Materials and Methods

### Study design

This study was designed to evaluate the involvement of PD-L2 both in in vivo models and human clinical specimens and to characterize structural, biochemical, functional and therapeutic efficacy of a sPD-1 mutants with ultra-high binding affinity to both PD-L1 and PD-L2. All tumor microarray specimens were commercially purchased from US Biomax (Derwood, MD). A board certified veterinarian pathologist performed the Immunohistochemical scoring of the stained tumor microarray specimens. Mouse in vivo studies were conducted under the approval of AAAPLAC at Stanford University and ChemPartner, Shanghai. Sample size for animal studies was based on previous experience with similar in vivo studies. All animals were randomly assigned to treatment groups. Samples were not excluded from studies except for animals that required early termination due to illness that is unrelated to the study. Endpoints of experiments were defined in advance for each experiment. Tumor growth curves were presented for studies where tumor growth was measurable and Kaplan-Meier curves were used for ID8 orthotropic ovarian cancer model. Appropriate statistical analysis was used for each experimental study.

### Cell lines

MC38, ID8, ID8-PDL1 CRISPR, B16/OVA, MC38-hPD-L1, MC38-hPD-L2, MC38-hPD-L2-PD-L1 CRISPR, Hep3B-OS8-hPD-L1, Hep3B-OS8-hPD-L2 were maintained in Dulbecco’s Modified Eagle’s Medium (DMEM) supplemented with 10% fetal bovine serum (FBS) and 1% antibiotics in a humidified 37 °C, 5% CO2 incubator. Cells were trypsinized and passaged at 80% confluency.

### Synthesis of yeast-displayed sPD-1 library

DNA encoding human PD-1 extracellular domain, amino acids Leu25 – Val170, was cloned into the pCT yeast display plasmid using *NheI* and *BamHI* restriction sites. Sequence numbering was done to match that used in Sasaki *et al*<u>^25^</u> to facilitate comparisons to their work with the wild-type proteins. An error-prone library was created using the PD-1 extracellular domain DNA as a template and mutations were introduced by using low-fidelity Taq polymerase (Invitrogen) and the nucleotide analogs 8-oxo-dGTP and dPTP (TriLink Biotech). Six separate PCR reactions were performed in which the concentration of analogs and the number of cycles were varied to obtain a range of mutation frequencies; five cycles (200 µM), ten cycles (2, 20, or 200 µM), and 20 cycles (2 or 20 µM). Products from these reactions were amplified using forward and reverse primers each with 50 bp homology to the pCT plasmid in the absence of nucleotide analogs. Amplified DNA was purified using gel electrophoresis and pCT plasmid was digested with *NheI* and *BamHI*. Purified mutant cDNA and linearized plasmid were electroporated in a 5:1 ratio by weight into EBY100 yeast where they were assembled *in vivo* through homologous recombination^40^. Library size was estimated to be 7.4×10^7^ by dilution plating.

### Library screening

Yeast displaying high affinity PD-1 mutants were isolated from the library using fluorescence-activated cell sorting (FACS). For FACS rounds 1 – 3, equilibrium binding sorts were performed in which yeast were incubated at room temperature in phosphate buffered saline with 0.1% BSA (PBSA) with the following nominal concentrations of PD-L1 (R&D Systems): sort 1) 10 nM PD-L1 for 3 h; sort 2) 2 nM PD-L1 for 3 h; sort 3) 0.2 nM PD-L1 for 24 h. After incubation with PD-L1, yeast were pelleted, washed, and resuspended in PBSA with 1:250 of chicken anti-c-Myc (Invitrogen) for 1 h at 4 °C. Yeast were then washed, pelleted and secondary labeling was performed for on ice for 30 min using PBSA with a 1:100 dilution of goat anti-chicken Alexa Fluor 555 (Invitrogen) and mouse anti-HIS Hilyte Fluor 488 (Anaspec).

For FACS rounds 4 – 6, kinetic off-rate sorts were conducted in which yeast were incubated with 2 nM PD-L1 for 3 hours at room temperature, after which cells were washed twice to remove excess unbound PD-L1, and resuspended in PBSA containing a ∼50 fold molar excess of PD-1 (R&D Systems) to render unbinding events irreversible. The length of the unbinding step was as follows: sort 4) 4 h; sort 5) 4 h; sort 6) 24 h, with all unbinding reactions performed at room temperature. During the last hour of the dissociation reaction, chicken anti-c-Myc was added to a final dilution of 1:250. Yeast were pelleted, washed, and secondary labeling was performed as previously described. Labeled yeast were sorted by FACS using a Vantage SE flow cytometer (Stanford FACS Core Facility) and CellQuest software (Becton Dickinson). Sorts were conducted such that the 1–3% of clones with the highest PD-L1 binding/c-Myc expression ratio were selected, enriching the library for clones with the highest binding affinity to PD-L1. In sort 1, 10^8^ cells were screened and subsequent rounds analyzed a minimum of ten-fold the number of clones collected in the prior sort round to ensure adequate sampling of the library diversity. Selected clones were propagated and subjected to further rounds of FACS. Following sorts 5 and 6, plasmid DNA was recovered using a Zymoprep kit (Zymo Research Corp.), transformed into XL-1 blue supercompetent cells, and isolated using plasmid miniprep kit (Qiagen). Sequencing was performed by Sequetech Corp.

Analysis of yeast-displayed sort products was performed using the same reagents and protocols and described for the library sorts. Samples were analyzed on a FACSCalibur (BD Biosciences) and data was analyzed using FlowJo software (Treestar Inc.).

### Recombinant protein production

sPD-1 wild-type and mutants linked to hIgG4Fc were cloned into the pCPC plasmid with signaling peptide (ChemPartner, Shanghai) and transiently transfected into HEK293 Human embryonic kidney cells and cultured in 4L OPM-293 CD03. Products were then purified through protein A resin MabSelect SuRe (GE Healthcare) and further isolated using superdex 200 size exclusion chromatography. Each purification steps were followed by SDS-PAGE confirmation and SEC-HPLC analysis. Final products were endotoxin tested and confirmed at 0.09EU/mg.

### Computational modeling analysis

Human PD1/PDL1 complex structure is available in Protein Data Bank, the PDBID is 4ZQK and 5IUS. However, in both of the PDB structures, some residues of PD1 are missing in a loop area, and some of the residues were mutated comparing with wild-type PD1. The missing loop and mutated the residue were reconstructed back to the wild-type PD1 based on crystal structure 4ZQK. The mutated PD1 structure was modeled using Residue Scanning module in Schrodinger Suite. The residues within 5.0 Å of the mutated residue were refined with side-chain prediction and backbone sampling. The calculation of variation of binding affinity and complex stability were also performed for multiple mutations. The more negative value of binding affinity or stability indicates more increase of them. The homology model of PD-L2 was built to investigate the interaction between human PD1 and PD-L2. After homology search in PDB non-redundant data set, three PDB structures of mouse PD-L2 were chosen to be templates. PD1 binding interface comparison between human PD-L2 model and mouse PD-L2 crystal structure (PDBID: 3BP5).

### Immunoassays

Slides were de-paraffined and antigen retrieval carried out using 10mM Citric Acid Buffer, 0.05% Tween 20, pH 6 Slides were removed from buffer and cooled at room temperature for 15 minutes. Quenched endogenous peroxidase with 1:10 dilution of 34% hydrogen peroxide and water for 15 minutes. Avidin and Biotin blocker were added for 15 minutes each. Protein block using 2% fetal calf serum was added for 20 minutes. The serum and antibodies were diluted in PBT (1XPBS, 0.1% BSA, 0.2%, 0.01% Tween 20). αHuman PD-L1 1:500 (#AF154, R&D systems,

Minneapolis, MN), αΗuman PD-L2 1:500 (#MABC1120, EMD Millipore, MA), αΗuman CD8 1:500 (#Ab4005, Abcam, Cambridge, MA), αMouse PD-L1 1:750 (#17952-1-AP, Proteintech, IL), αMouse PD-L2 1:750 (#Ab21107, Abcam, Cambridge, MA) and αΜouse CD8 1:500 (#Ab203035, Abcam, Cambridge, MA) antibodies incubated overnight at °C. For immunohistochemistry, anti-rabbit and anti-rat were added on each slide and incubated at 37°C for 30 minutes then incubated with STREP-HRP for 30 minutes at 37°C and signals developed using DAB substrate kit (#34002, ThermoFisher Scientific, Waltham, MA). For immunofluorescence detection, all staining were carried out in dark, Secondary antibody1:400 diluted in 2% BSA and 0.1% tween 20 in PBS was incubated at room temperature for 1 hour. Sudan Black B 0.2% solution in 70% ethanol through syringe filter, pre-warmed to RT was added to the slides for 10 minutes at room temperature. DAPI (#F6057, Sigma-Aldrich, St. Louis, MO) 0.1µg/mL was used for counterstain and coverslip applied. ELISA quantification of human IgG4 (#BMS2059, ThermoFisher Scientific, Waltham, MA) was carried out per manufacturer’s instruction.

### Surface Plasmon Resonance

Human PD-L1 (#10084-H08H, SinoBiological, Wayne, PA) and human PD-L2 (#10292-H08H, SinoBiological, Wayne, PA) were used as analyte. Running buffer HBS-EP+ (10mM HEPES, 150mM NaCl, 3mM EDTA and 0.05% P20, pH 7.4) was prepared. Flow rate was performed at 30 µL/min. Immobilization of anti-human Fc IgG on CM5 sensorchip. PD1-Fc was captured to the immobilized sensorchip. An injection of serial diluted analyte was used. Glycine pH 1.5 for 30s was used as the surface regeneration. Multiple cycle kinetics was carried out on Biacore T200. Kd analysis calculated using the Biacore T200 Evaluation Software (GE Healthcare Life Sciences).

### Flow Cytometry Based Binding Analysis

The MC38-hPD-L1, MC38-hPD-L2, Hep3B-OS8-hPD-L1, Hep3B-OS8-hPD-L2 and parental MC38 cells were cultured in standard tissue culture condition. Cells were harvested and the supernatant discarded then dispensed onto a staining plate at 3x10^5^ cells per well. The plate was centrifuged at 300g at 4°C for 5 minutes. Various concentrations of sPD-1 mutants and negative control were diluted in FACS buffer containing 2% FBS, 100µL/well was added. Cells were incubated for 1 hour at 4°C and washed twice with 200µL FACS buffer and centrifuged at 300g for 5 minutes. The supernatant was discarded before and after each wash. Cells were re-suspended at 100µL/well with 1:1000 diluent with anti-human IgG-Alexa 488 (#A28175, ThermoFisher, Waltham, MA). Plates were incubated for 1 hour at 4°C. Cells were washed twice with FACS buffer and centrifuged at 300g for 5 minutes. Supernatant was discarded and cells were re-suspended in 100µL cold PBS. The cells were kept in the dark and FACS analysis carried out on FACS CantoII, (BD Biosciences, San Jose, CA).

### Receptor Blocking Assay

Hep3B-OS8-hPD-L1, Hep3B-OS8-hPD-L2, MC38-hPD-L1 and MC38-hPD-L2 cells were cultured in PRMI 1640 medium, 10% FBS, G418 and hygro, in T175 flask to a confluency of 60-80%. The cells were harvested and re-suspended into dilution buffer. cells were dispensed into round bottom 96-well plate corning #3799 at the density of 2x10^5^ cells/well. sPD-1 mutant at multiple concentration: 600, 120, 24, 4.8, 0.96, 0.192, 0.00384 nM and 0 nM were re-suspended in dilution Buffer. sPD-1 wild-type were also prepared in dilution Buffer. The plate was centrifuged at 500g for 3 minutes and supernatants discarded. Cells were re-suspended with 50µL antibody followed by adding 50µL of ligand. The plates were incubated at 4°C for 2 hours followed by centrifugation and washed. Streptavidin-Alexa Flour 488 (#S11223, Thermo Fisher, Waltham, MA) diluted 1:1000 in dilution Buffer was prepare and incubated with cells at 4°C for 1 hour. The plate was centrifuged and washed then re-suspended in 200µL FACS buffer. Cells ran on FACS Canto II (BD Biosciences, San Jose, CA). Curves with MFI values were drawn and IC50s and tops were calculated.

### T Cell Stimulation and T proliferation Assay

Blood sample from healthy donors were diluted by equal volume of sterile PBS and mixed by gentle shaking. 15mL of Ficoll-Paque PLUS (10-1440-02, GE Healthcare, Pittsburgh, PA) medium was transferred into a fresh 50mL centrifuge tube with a Ficoll and blood volume ration of 3:4. The diluted blood sample was then carefully layered onto the surface of the Ficoll medium to avoid mixing. The tube was centrifuged at 400 g for 30 minutes at 20°C with the max acceleration and min deceleration settings during the centrifugation. 4 interfaces was observed after the centrifugation with layers of plasma, mononuclear cells, Ficoll medium, and RBCs seen from top to bottom. Gently removing the layer of mononuclear cells were carefully transferred into a new sterile centrifuge tube. Sterile PBS buffer was added into the collected PBMCs for washing. The tube was centrifuged at 300xg for 10 minutes at 20°C with the max acceleration and min deceleration settings during the centrifugation. Cells were re-suspended with 10%FBS+RPMI 1640 for assay. Mycoplasma-free Hep3b-hPDL1-OS8 cells were prepared in a 15cm dish with the confluency kept at 60-80% before use. Cells were trypsinized with TrypLE^TM^ Express, (#12605-036, Thermo Fisher, Waltham, MA) were collected to a 50mL centrifuge tube and centrifuged at 1000rpm for 5 minutes. The supernatant was discarded and cells were re-suspended with mitomycin (#H33020786, Zhejiang, China) 10µg/mL in 5-10mL 10%FBS + RPMI 1640 medium. Cells were incubated at 37°C for an hour. After incubation, cells were washed in PBS cell counting with cytometer. Cells were centrifuged at 1000rpm for 5 minutes at 20°C. Cells were then re-suspended with 10%FBS + RPMI 1640 assay. PBMC stimulation was performed by adding PBMC at 5E4 cell/well with 100µL/well on a 96 well microplate. 50 µL/well of a series of 5X OD1 v2, 5X PD1 WT, or hIgG4 protein solution was added followed by APC Hep3b-hPDL1-OS8 at 5000 cell/well for 50µL/well. The mixed solution was incubated at 37°C for 72 hours. The supernatant was harvested and placed in -20°C before ELISA test. The IFN-γ concentration was tested with the ELISA kit. Quantification of IFN-γ was determined by quantikine ELISA kits purchased from R&D. The assay procedure outlined by the Human IFN-γ ELISA kit, R&D, Cat#DY285 was followed. The plate was read with the ELISA plate reader at 450nm wavelength.

### Expression of PD-L2 in Lentiviral Transduced Hep3B-OS8 Cells and MC38 Cells

Hep3B-OS8(4E8) cell line was transduced with pLVX-IRES-hygro-hPDL2:CP in house, raising in 1640 medium. MC38 cells were transduced with pLVX-IRES-hygro-hPD-L2. Cells were harvested according to standard procedures and the supernatant was discarded. Cells were dispensed into a round bottom 96-well staining plate: corning, REF#3799 with 3e5 cells per well. The plate was placed in the centrifuge at 300g at 4°C for 5 minutes. 100µL of rabbit anti-hPD-L2 in titrated 1xPBS+2%FBS FACS buffer was added to each well and incubated for one hour at 4°C. Cells were washed twice with 200µL FACS buffer and centrifuged at 300g for 5 minutes. Supernatant was discarded before and after each wash. Cells were re-suspended at 100µL/well with 1:1000 dilution of secondary donkey-anti-rabbit IgG(H+L) Alexa488 life technology antibody and incubated for 1 hour at 4°C. Cells were again washed twice with 200µL FACS buffer and centrifuged at 300g for 5 minutes. Supernatant was discarded before and after each wash. Cells were re-suspended in 100µL PBS. Cells were kept in the dark and submitted for FACS analysis.

### In vivo tumor studies

All animal experiments were reviewed and approved by the Institutional Animal Care and Use Committee (IACUC) at Stanford University and animal ethical committee at ChemPartner, Shanghai. C57B/6 mice with hPD-L1 knock-in were purchased from Biocytogen (Beijing, China) and wild-type C57/B6 mice were purchased from Biocytogen (Beijing, China) or The Jackson Laboratory (Bar Harbor, Maine). PD-L1 knockout mice on C57B/6 background were kindly gifted by (Dr. Dean W. Felsher at Stanford University). Female mice age 6-8 weeks were used for ID8 ovarian tumor studies and male mice aged 6-8 weeks were used for MC38 colorectal studies and B16/OVA melanoma studies. Mice were housed in pathogen-free animal facility, kept under constant temperature and humidity and controlled 12 h light-dark cycles. For MC38, MC38-hPD-L1, MC38-hPD-L2 colorectal study, and B16/OVA melanoma study, 1x10^6 cells were injected subcutaneously and randomly assigned to control or appropriate treatment group upon confirmation of tumor growth approximately 7 days post tumor inoculation. Tumor growth was measured throughout the study and body weight recorded. Animals were sacrificed at the same time when majority of tumors reached ethical endpoint. For ID8 ovarian study, 5x10^6 cells were injected intraperitoneal, and each animal terminated upon development of peritoneal ascite for survival analysis. For tumor studies conducted in PD-L1 knockout mice, 10x10^6 ID8 cells suspended in 50% Matrigel (#356230 Corning, Cornng, NY) were injected subcutaneously.

### Statistical analysis

The Pearson correlation was used for all correlation analysis of tumor specimens. IHC H score values were determined by a board certified veterinarian pathologist. All tumor volume, survival, quantification of *in vivo* immunohistochemistry were conducted using GraphPad Prism software (GraphPad Software Inc). ANOVA with Tukey-Kramer test was used for comparing multiple treatment groups with each other. *P* < 0.05 was considered significant. Repeated measure ANOVA was used for comparing multiple treatment groups measured over time. Statistical analysis of survival curves was conducted for the ID8 survival studies. A log-rank (Mantel-Cox) test was performed to compare mean survival among groups; *P* ≤ 0.05 was considered statistically significant.

## Acknowledgments

This work was supported by NIH grants CA67166, CA197713, and CA198291, the Silicon Valley Foundation, the Sydney Frank Foundation, and the Kimmelman Fund (A.J.G.). Authors like to thank Hannah Elizabeth Skolnik for technical support.

## Reference

1. Herbst RS, Baas P, Kim DW, Felip E, Perez-Gracia JL, Han JY, Molina J, Kim JH, Arvis CD, Ahn MJ, et al. Pembrolizumab versus docetaxel for previously treated, PD-L1-positive, advanced non-small-cell lung cancer (KEYNOTE-010): a randomised controlled trial. Lancet. 2016;387(10027):1540–50.

2. Larkin J, Minor D, D’Angelo S, Neyns B, Smylie M, Miller WH, Jr., Gutzmer R, Linette G, Chmielowski B, Lao CD, et al. Overall Survival in Patients With Advanced Melanoma Who Received Nivolumab Versus Investigator’s Choice Chemotherapy in CheckMate 037: A Randomized, Controlled, Open-Label Phase III Trial. J Clin Oncol. 2018;36(4):383–90.

3. Schachter J, Ribas A, Long GV, Arance A, Grob JJ, Mortier L, Daud A, Carlino MS, McNeil C, Lotem M, et al. Pembrolizumab versus ipilimumab for advanced melanoma: final overall survival results of a multicentre, randomised, open-label phase 3 study (KEYNOTE-006). Lancet. 2017;390(10105):1853–62.

4. Wang C, Thudium KB, Han M, Wang XT, Huang H, Feingersh D, Garcia C, Wu Y, Kuhne M, Srinivasan M, et al. In vitro characterization of the anti-PD-1 antibody nivolumab, BMS-936558, and in vivo toxicology in non-human primates. Cancer Immunol Res. 2014;2(9):846–56.

5. Rodriguez CP, Wu Q, Voutsinas J, Fromm JR, Jiang X, Pillarisetty VG, Lee SM, Santana-Davila R, Goulart B, Baik CS, et al. A phase II trial of pembrolizumab and vorinostat in recurrent metastatic head and neck squamous cell carcinomas and salivary gland cancer. Clin Cancer Res. 2019.

6. Varga A, Piha-Paul S, Ott PA, Mehnert JM, Berton-Rigaud D, Morosky A, Yang P, Ruman J, and Matei D. Pembrolizumab in patients with programmed death ligand 1-positive advanced ovarian cancer: Analysis of KEYNOTE-028. Gynecol Oncol. 2019;152(2):243–50.

7. Wang BC, Zhang ZJ, Fu C, and Wang C. Efficacy and safety of anti-PD-1/PD-L1 agents vs chemotherapy in patients with gastric or gastroesophageal junction cancer: a systematic review and meta-analysis. Medicine (Baltimore). 2019;98(47):e18054.

8. Imai Y, Hasegawa K, Matsushita H, Fujieda N, Sato S, Miyagi E, Kakimi K, and Fujiwara K. Expression of multiple immune checkpoint molecules on T cells in malignant ascites from epithelial ovarian carcinoma. Oncol Lett. 2018;15(5):6457–68.

9. Pietzner K, Nasser S, Alavi S, Darb-Esfahani S, Passler M, Muallem MZ, and Sehouli J. Checkpoint-inhibition in ovarian cancer: rising star or just a dream? J Gynecol Oncol. 2018;29(6):e93.

10. Tran L, Allen CT, Xiao R, Moore E, Davis R, Park SJ, Spielbauer K, Van Waes C, and Schmitt NC. Cisplatin Alters Antitumor Immunity and Synergizes with PD-1/PD-L1 Inhibition in Head and Neck Squamous Cell Carcinoma. Cancer Immunol Res. 2017;5(12):1141–51.

11. Rini BI, Plimack ER, Stus V, Gafanov R, Hawkins R, Nosov D, Pouliot F, Alekseev B, Soulieres D, Melichar B, et al. Pembrolizumab plus Axitinib versus Sunitinib for Advanced Renal-Cell Carcinoma. N Engl J Med. 2019;380(12):1116–27.

12. Tree AC, Jones K, Hafeez S, Sharabiani MTA, Harrington KJ, Lalondrelle S, Ahmed M, and Huddart RA. Dose-limiting Urinary Toxicity With Pembrolizumab Combined With Weekly Hypofractionated Radiation Therapy in Bladder Cancer. Int J Radiat Oncol Biol Phys. 2018;101(5):1168–71.

13. Zhang L, Conejo-Garcia JR, Katsaros D, Gimotty PA, Massobrio M, Regnani G, Makrigiannakis A, Gray H, Schlienger K, Liebman MN, et al. Intratumoral T cells, recurrence, and survival in epithelial ovarian cancer. N Engl J Med. 2003;348(3):203–13.

14. Luo Z, Wang Q, Lau WB, Lau B, Xu L, Zhao L, Yang H, Feng M, Xuan Y, Yang Y, et al. Tumor microenvironment: The culprit for ovarian cancer metastasis? Cancer Lett. 2016;377(2):174–82.

15. Ishida M, Iwai Y, Tanaka Y, Okazaki T, Freeman GJ, Minato N, and Honjo T. Differential expression of PD-L1 and PD-L2, ligands for an inhibitory receptor PD-1, in the cells of lymphohematopoietic tissues. Immunol Lett. 2002;84(1):57–62.

16. Latchman Y, Wood CR, Chernova T, Chaudhary D, Borde M, Chernova I, Iwai Y, Long AJ, Brown JA, Nunes R, et al. PD-L2 is a second ligand for PD-1 and inhibits T cell activation. Nat Immunol. 2001;2(3):261–8.

17. Lesterhuis WJ, Punt CJ, Hato SV, Eleveld-Trancikova D, Jansen BJ, Nierkens S, Schreibelt G, de Boer A, Van Herpen CM, Kaanders JH, et al. Platinum-based drugs disrupt STAT6-mediated suppression of immune responses against cancer in humans and mice. J Clin Invest. 2011;121(8):3100–8.

18. Yamazaki T, Akiba H, Iwai H, Matsuda H, Aoki M, Tanno Y, Shin T, Tsuchiya H, Pardoll DM, Okumura K, et al. Expression of programmed death 1 ligands by murine T cells and APC. J Immunol. 2002;169(10):5538–45.

19. Liang SC, Latchman YE, Buhlmann JE, Tomczak MF, Horwitz BH, Freeman GJ, and Sharpe AH. Regulation of PD-1, PD-L1, and PD-L2 expression during normal and autoimmune responses. Eur J Immunol. 2003;33(10):2706–16.

20. Lazar-Molnar E, Yan Q, Cao E, Ramagopal U, Nathenson SG, and Almo SC. Crystal structure of the complex between programmed death-1 (PD-1) and its ligand PD-L2. Proc Natl Acad Sci U S A. 2008;105(30):10483–8.

21. Youngnak P, Kozono Y, Kozono H, Iwai H, Otsuki N, Jin H, Omura K, Yagita H, Pardoll DM, Chen L, et al. Differential binding properties of B7-H1 and B7-DC to programmed death-1. Biochem Biophys Res Commun. 2003;307(3):672–7.

22. Derks S, Nason KS, Liao X, Stachler MD, Liu KX, Liu JB, Sicinska E, Goldberg MS, Freeman GJ, Rodig SJ, et al. Epithelial PD-L2 Expression Marks Barrett’s Esophagus and Esophageal Adenocarcinoma. Cancer Immunol Res. 2015;3(10):1123–9.

23. Pujade-Lauraine E, Fujiwara K, Dychter SS, Devgan G, and Monk BJ. Avelumab (anti-PD-L1) in platinum-resistant/refractory ovarian cancer: JAVELIN Ovarian 200 Phase III study design. Future Oncol. 2018;14(21):2103–13.

24. de Groot J, Penas-Prado M, Alfaro-Munoz KD, Hunter K, Pei B, O’Brien B, Weathers SP, Loghin M, Kamiya Matsouka C, Yung WKA, et al. Window-of-opportunity clinical trial of pembrolizumab in patients with recurrent glioblastoma reveals predominance of immune-suppressive macrophages. Neuro Oncol. 2019.

25. Inc. P. Avelumab in Previously Untreated Patients With Epithelial Ovarian Cancer (JAVELIN OVARIAN 100).

26. Pfizer ESa. EMD Serono and Pfizer Provide Update on Phase III JAVELIN Gastric 100 Trial. https://w.pfizer.com/news/press-release/press-release-detail/emd_serono_and_pfizer_provide_update_on_phase_iii_javelin_gastric_100_trial. Accessed November, 8th, 2019.

27. Fradet Y, Bellmunt J, Vaughn DJ, Lee JL, Fong L, Vogelzang NJ, Climent MA, Petrylak DP, Choueiri TK, Necchi A, et al. Randomized phase III KEYNOTE-045 trial of pembrolizumab versus paclitaxel, docetaxel, or vinflunine in recurrent advanced urothelial cancer: results of > 2 years of follow-up. Ann Oncol. 2019.

28. Zhang F, Qi X, Wang X, Wei D, Wu J, Feng L, Cai H, Wang Y, Zeng N, Xu T, et al. Structural basis of the therapeutic anti-PD-L1 antibody atezolizumab. Oncotarget. 2017;8(52):90215–24.

29. Kariolis MS, Miao YR, Jones DS, 2nd, Kapur S, Mathews, II, Giaccia AJ, and Cochran JR. An engineered Axl ’decoy receptor’ effectively silences the Gas6-Axl signaling axis. Nat Chem Biol. 2014;10(11):977–83.

30. Kariolis MS, Miao YR, Diep A, Nash SE, Olcina MM, Jiang D, Jones DS, 2nd, Kapur S, Mathews, II, Koong AC, et al. Inhibition of the GAS6/AXL pathway augments the efficacy of chemotherapies. J Clin Invest. 2017;127(1):183–98.

31. Silva JP, Vetterlein O, Jose J, Peters S, and Kirby H. The S228P mutation prevents in vivo and in vitro IgG4 Fab-arm exchange as demonstrated using a combination of novel quantitative immunoassays and physiological matrix preparation. J Biol Chem. 2015;290(9):5462–9.

32. Tang S, and Kim PS. A high-affinity human PD-1/PD-L2 complex informs avenues for small-molecule immune checkpoint drug discovery. Proc Natl Acad Sci U S A. 2019;116(49):24500–6.

33. Burova E, Hermann A, Waite J, Potocky T, Lai V, Hong S, Liu M, Allbritton O, Woodruff A, Wu Q, et al. Characterization of the Anti-PD-1 Antibody REGN2810 and Its Antitumor Activity in Human PD-1 Knock-In Mice. Mol Cancer Ther. 2017;16(5):861–70.

34. Moehler M, Ryu MH, Dvorkin M, Lee KW, Coskun HS, Wong R, Chung HC, Poltoratsky A, Tsuji A, Yen CJ, et al. Maintenance avelumab versus continuation of first-line chemotherapy in gastric cancer: JAVELIN Gastric 100 study design. Future Oncol. 2019;15(6):567–77.

35. Kudo T, Hamamoto Y, Kato K, Ura T, Kojima T, Tsushima T, Hironaka S, Hara H, Satoh T, Iwasa S, et al. Nivolumab treatment for oesophageal squamous-cell carcinoma: an open-label, multicentre, phase 2 trial. Lancet Oncol. 2017;18(5):631–9.

36. Nazareth MR, Broderick L, Simpson-Abelson MR, Kelleher RJ, Jr., Yokota SJ, and Bankert RB. Characterization of human lung tumor-associated fibroblasts and their ability to modulate the activation of tumor-associated T cells. J Immunol. 2007;178(9):5552–62.

37. Messal N, Serriari NE, Pastor S, Nunes JA, and Olive D. PD-L2 is expressed on activated human T cells and regulates their function. Mol Immunol. 2011;48(15-16):2214–9.

38. (NCI) NCI. AMP-224, a PD-1 Inhibitor, With Stereotactic Body Radiation Therapy in Metastatic Colorectal Cancer.

39. Huber S, Hoffmann R, Muskens F, and Voehringer D. Alternatively activated macrophages inhibit T-cell proliferation by Stat6-dependent expression of PD-L2. Blood. 2010;116(17):3311–20.

40. Lesterhuis WJ, Steer H, and Lake RA. PD-L2 is predominantly expressed by Th2 cells. Mol Immunol. 2011;49(1-2):1–3.

41. Yearley JH, Gibson C, Yu N, Moon C, Murphy E, Juco J, Lunceford J, Cheng J, Chow LQM, Seiwert TY, et al. PD-L2 Expression in Human Tumors: Relevance to Anti-PD-1 Therapy in Cancer. Clin Cancer Res. 2017;23(12):3158–67.

42. Hebenstreit D, Wirnsberger G, Horejs-Hoeck J, and Duschl A. Signaling mechanisms, interaction partners, and target genes of STAT6. Cytokine Growth Factor Rev. 2006;17(3):173–88.

43. Li Y, Liang Z, Tian Y, Cai W, Weng Z, Chen L, Zhang H, Bao Y, Zheng H, Zeng S, et al. High-affinity PD-1 molecules deliver improved interaction with PD-L1 and PD-L2. Cancer Sci. 2018;109(8):2435–45.

44. Maute RL, Gordon SR, Mayer AT, McCracken MN, Natarajan A, Ring NG, Kimura R, Tsai JM, Manglik A, Kruse AC, et al. Engineering high-affinity PD-1 variants for optimized immunotherapy and immuno-PET imaging. Proc Natl Acad Sci U S A. 2015;112(47):E6506–14.

45. Deng R, Bumbaca D, Pastuskovas CV, Boswell CA, West D, Cowan KJ, Chiu H, McBride J, Johnson C, Xin Y, et al. Preclinical pharmacokinetics, pharmacodynamics, tissue distribution, and tumor penetration of anti-PD-L1 monoclonal antibody, an immune checkpoint inhibitor. MAbs. 2016;8(3):593–603.

46. Lin H, Wei S, Hurt EM, Green MD, Zhao L, Vatan L, Szeliga W, Herbst R, Harms PW, Fecher LA, et al. Host expression of PD-L1 determines efficacy of PD-L1 pathway blockade-mediated tumor regression. J Clin Invest. 2018;128(2):805–15.

