## Supplemental figure 1 for "Neutralizing PD-L1 and PD-L2 Enhances the Efficacy of Immune Checkpoint Inhibitors in Ovarian Cancer"

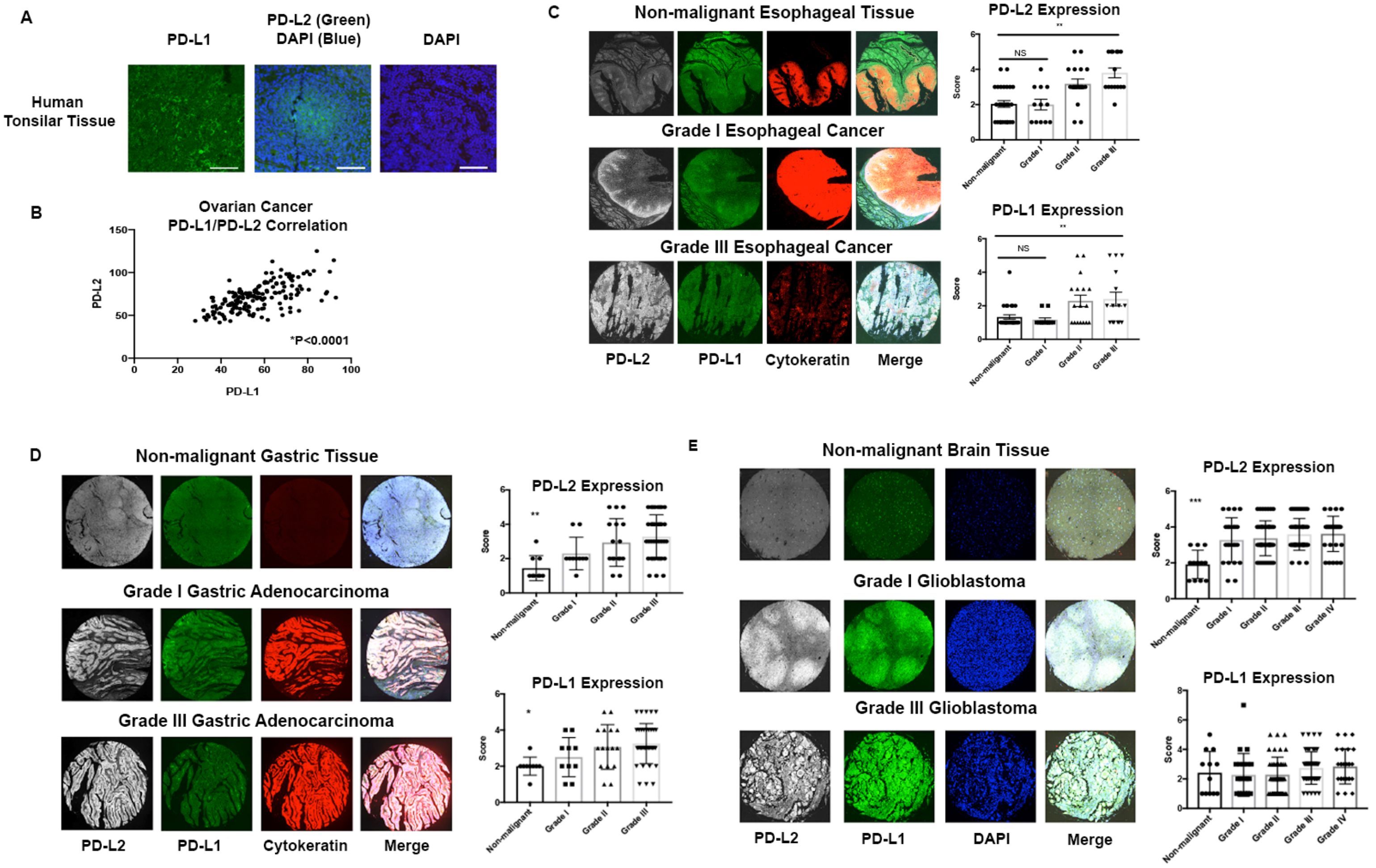


**Fig. S1. PD-L1 and PD-L2 expression analyzed in gastric, esophageal and brain cancer.** (A) IHC staining of human PD-L1 and PD-L2 on tonsil tissue showing distinct staining pattern for PD-L1 and PD-L2 expression. (B) Pearson correlation plot of PD-L1 and PD-L2 expression in ovarian cancer. (C) Representative images of gastric cancer tissue microarray containing both normal and malignant samples (N=76) were stained with anti-human PD-L2 (gray), anti-human PD-L1 (green) and cytokeratin (red) by fluorescent immunohistochemistry. The intensity of PD-L2 (right upper) and PD-L1 (right bottom) was scored and stratified according to the tumor grade. (D) Representative images of esophageal cancer tissue microarray containing both normal and malignant samples (N=72) were stained with anti-human PD-L2 (gray), anti-human PD-L1 (green) and cytokeratin (red) by fluorescent immunohistochemistry. The intensity of PD-L2 (right upper) and PD-L1 (right bottom) was scored and stratified according to the tumor grade. (E) Representative images of glioblastoma tissue microarray containing both normal and malignant samples (N=152) were stained with anti-human PD-L2 (gray), anti-human PD-L1 (green) and DAPI (blue) by fluorescent immunohistochemistry. The intensity of PD-L2 (right upper) and PD-L1 (right bottom) was scored and stratified according to the tumor grade.
