## Supplemental figure 2 to 15 for "Neutralizing PD-L1 and PD-L2 Enhances the Efficacy of Immune Checkpoint Inhibitors in Ovarian Cancer"

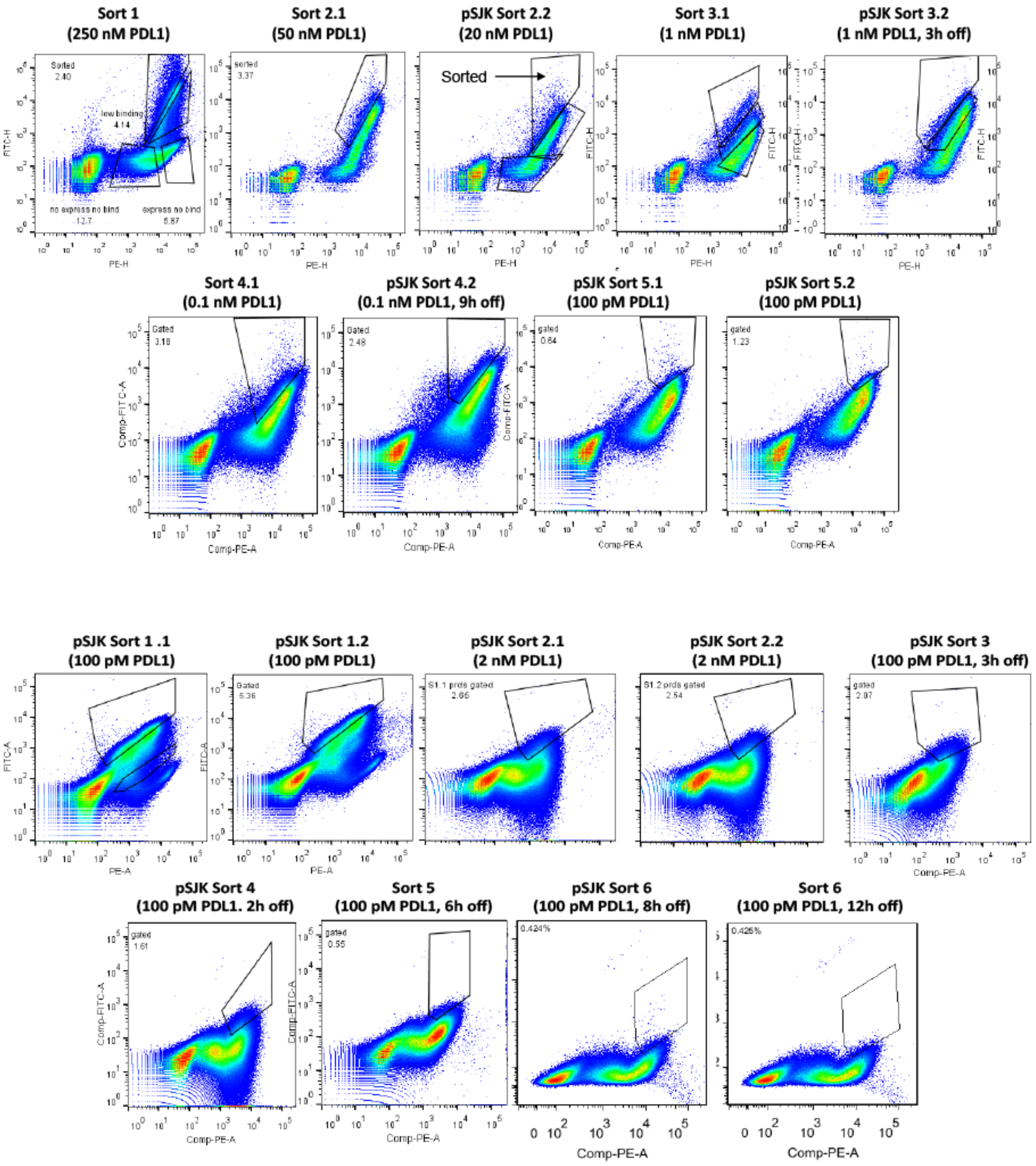


**Fig. S2. Directed evolution based sorting of mutant clones exhibiting superior binding characteristics.** The first two sorts were performed by decreasing the amount of PD-L1 incubated with the library in a sequential manner. Sort three to sort six was screened based on a combination of both decreasing the concentration of the ligand and kinetics off-rate sorting strategy in which clones were isolated based on the ability to bind PD-L1 in the presence of lower ligand concentration and longer incubation time in the presence of competitors. Percentages in each panel correspond to the gated subpopulation collected.

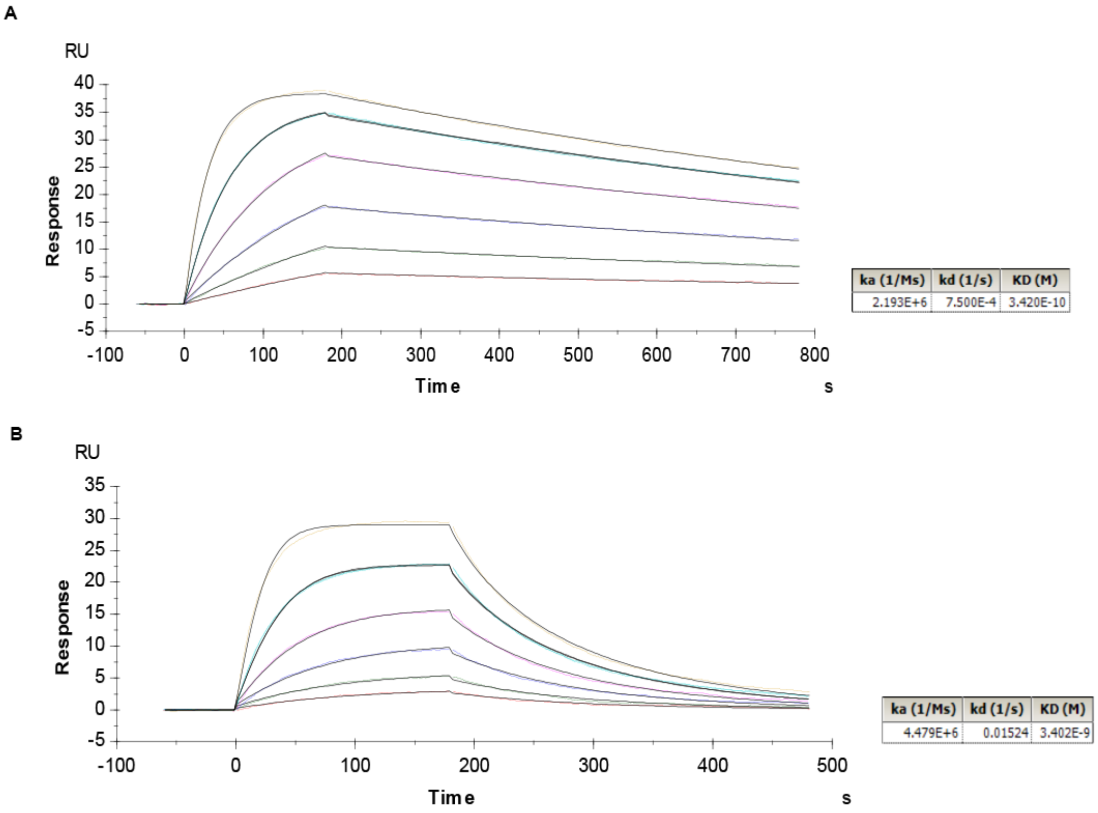


**Fig. S3. Analysis of sPD-1 mutant V2 binding kinetics to PD-L1 and PD-L2.** Binding analysis of sPD-1 mutant version 2 binding to kinetics to PD-L1 (A) and PD-L2 (b) by BIAcore T200 at 25 °C.

A

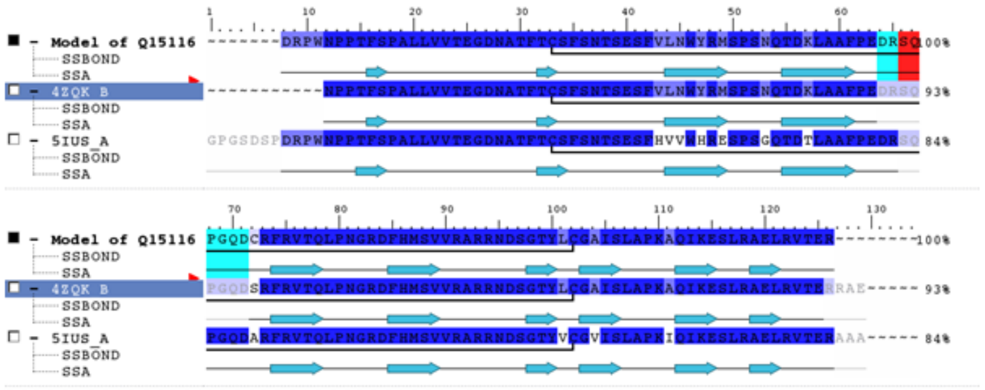


B

| Site in PD-1 | Surface Complementarity | | Hydrogen Bond with PD-L1 | |
|---|---|---|---|---|
| | WT | Single mutation | WT | Single mutation |
| G124S | 0.93 | 0.75 | 0 | 1 to Y123 |
| S127V | 0.89 | 0.64 | 0 | 0 |
| A132I | 0.72 | 0.85 | 1 to Q66 | 2 to Q66 |

**Fig. S4. Sequence alignment of human PD-1 and protein interaction between sPD-1 mutant and PD-L1.** (A) Sequence alignment of modeled wild-type PD-1 (Q15116) with PDB No. 4ZQK and 5IUS. The identical residues are marked in blue, the residues missing in both 4ZQK and 5IUS are highlighted in red and the residues missing only in 4ZQK are marked in cyan. (B) Analysis of surface complementarity and hydrogen bond for each of the three mutations within the binding interface on sPD-1 mutant when in complex with human PD-L1.

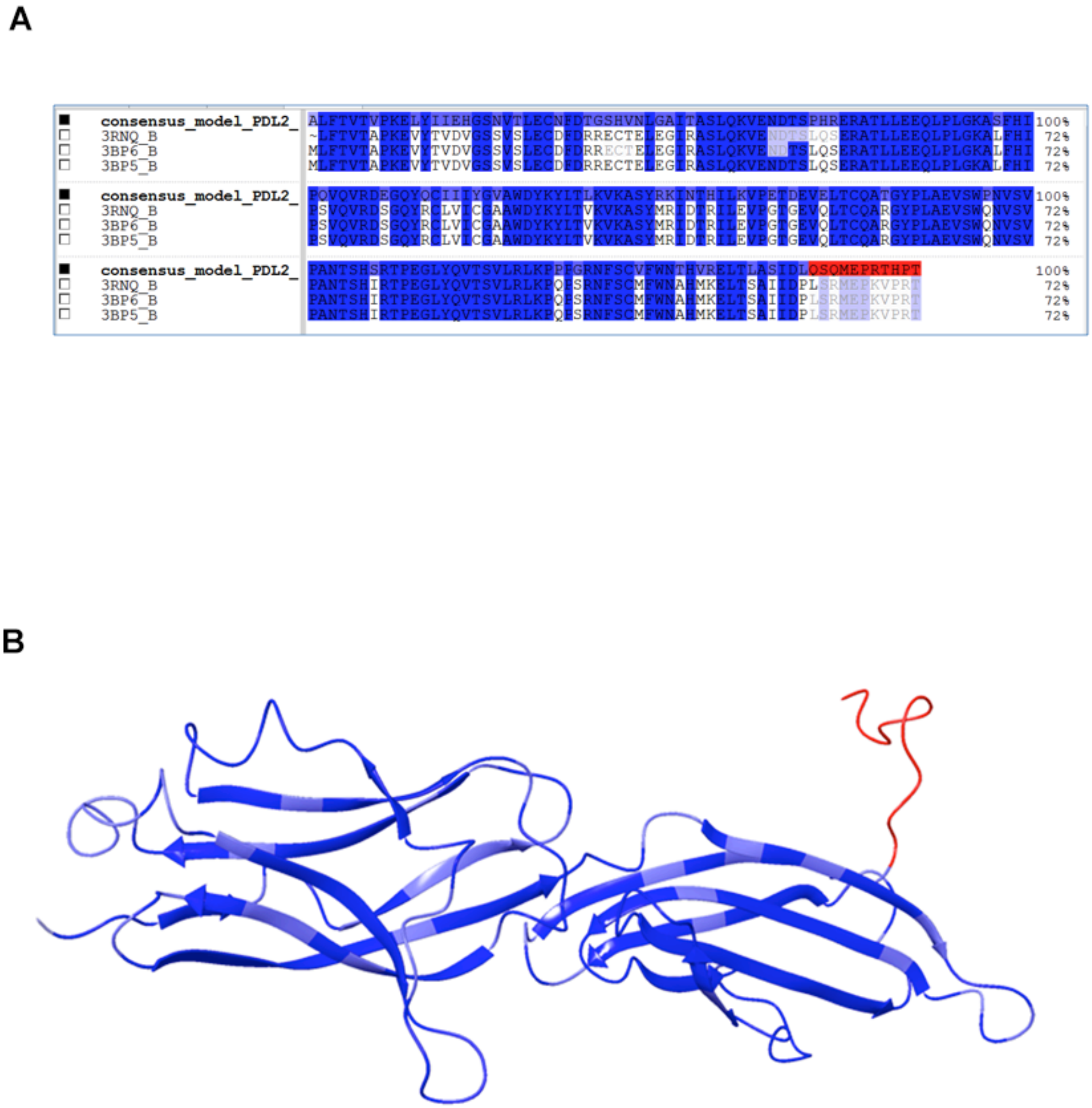


**Fig. S5. Modeling of human PD-L2 from mouse PD-L2.** (A) Sequence alignment betweenmodeled human PD-L2 and three mouse PD-L2 structures (PDB No. 3RNQ, 3BP6, 3BP5). The residues in missing loop are marked red. (B) Consensus model of human PD-L1.

**A**

| Site in PD1 | Surface Complementarity | | Hydrogen bond with PD-L2 | |
|---|---|---|---|---|
| | WT | Single mutation | WT | Single mutation |
| G124S | 0.80 | 0.85 | 0 | 0 |
| A132I | 0.83 | 0.86 | 1 to Q60 | 1 to Q60 |

**B**

| Mutations | d Affinity | d Stability (solvated) |
|---|---|---|
| 8 mutations | -10.62 | -10.7 |
| 7 mutations (N116S excluded ) | -10.21 | -12.19 |
| 4 outside mutations | 0.59 | -10.03 |
| 3 interface mutations | -11.69 | 8.59 |
| 2 mutations on missing loop | 0.01 | -4.95 |

**Fig. S6. Comparison of protein interaction between wild-type and mutated human PD-1/PD-L2 model.** (A) Analysis of surface complementarity and hydrogen bond for 2 mutations within the binding interface on sPD-1 mutant in co-complex with hPD-L2. (B) Calculated changes in binding affinity and protein stability (in kcal/mol) between sPD-1 mutant and hPD-L2 with mutations within and outside of the binding interface.

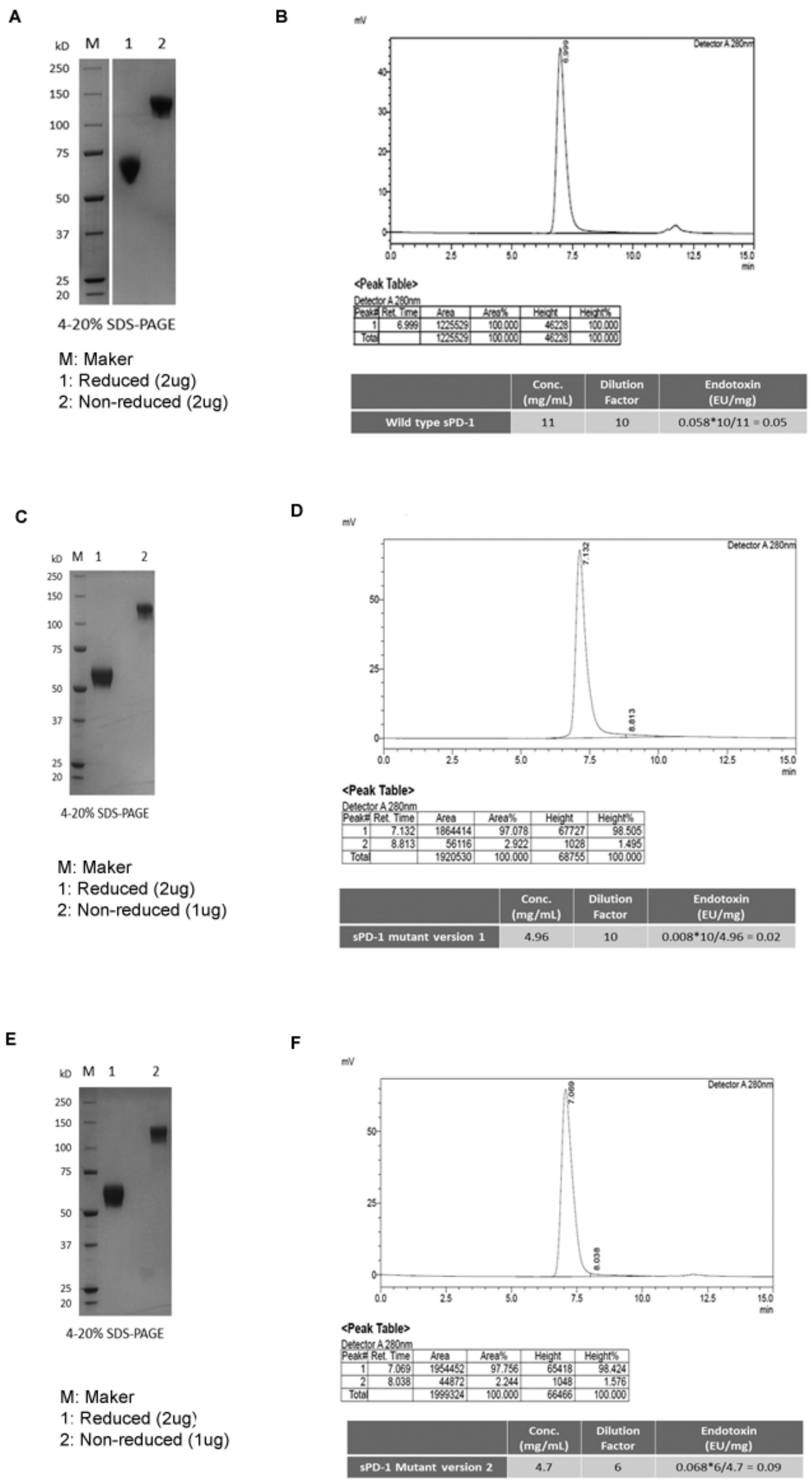


**Fig. S7. Recombinant protein production of sPD-1 mutants.** (A) SDS-PAGE of purified wild-type sPD-1 Fc, lane 1 reduced and lane 2 non-reduced. (B) SEC-HPLC analysis of purified wild-type sPD-1 Fc and endotoxin levels shown on the table below. (C) SDS-PAGE of purified sPD-1 mutant version 1, lane 1 reduced and lane 2 non-reduced. (D) SEC-HPLC analysis of purified sPD-1 mutant version 1 and endotoxin levels shown on the table below. (E) SDS-PAGE of purified sPD-1 mutant version 2, lane 1 reduced and lane 2 non-reduced. (F) SEC-HPLC analysis of purified sPD-1 mutant version 2 and endotoxin levels shown on the table below.

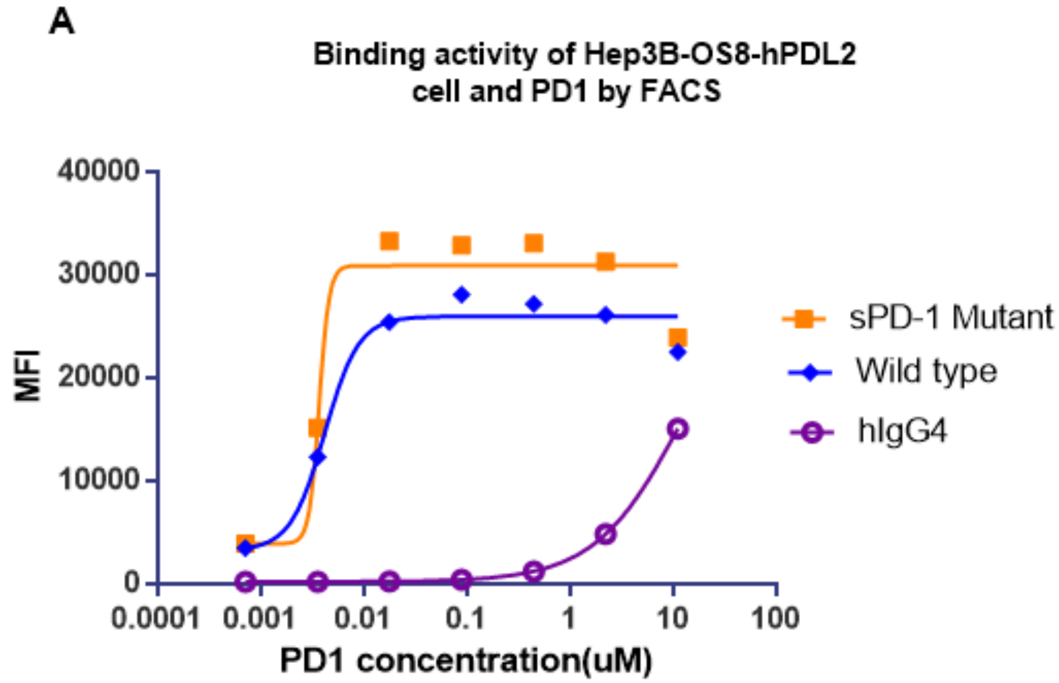


**Fig. S8**. **sPD-1 binding to human PD-L2.** Binding kinetics between sPD-1 mutant V2 binding to PD-L2 expressing Hep3B-OS8-PDL2 cells.

**A**

| | | cells\|Freq. of Parent (%) | cells\|Mean (Alexa Fluor 488-A) | coresponding specimen |
|---|---|---|---|---|
| 1E6/well during transduction | 4F3 | 86 | 4973.6 | Specimen_001_A1_A01_001.fcs |
| | 4G10 | 95.2 | 4730.4 | Specimen_001_A2_A02_002.fcs |
| | 4C3 | 94 | 4259.3 | Specimen_001_A3_A03_003.fcs |
| | 5 E9 | 97.2 | 4322.7 | Specimen_001_A4_A04_004.fcs |
| | 4C7 | 98.2 | 5820.7 | Specimen_001_A5_A05_005.fcs |
| | 4F11 | 65 | 3711.2 | Specimen_001_A6_A06_006.fcs |
| | 4F12 | 96.2 | 7110.8 | Specimen_001_A7_A07_007.fcs |
| | 4 E8 | 97.2 | 6691.2 | Specimen_001_A8_A08_008.fcs |
| 5E5/well during transduction | 5B11 | 95.1 | 7515 | Specimen_001_B1_B01_011.fcs |
| | 5C10 | 97.9 | 6117.9 | Specimen_001_B2_B02_012.fcs |
| | 4H7 | 99.7 | 11616.6 | Specimen_001_B3_B03_013.fcs |
| | 4B6 | 95.7 | 5717.5 | Specimen_001_B4_B04_014.fcs |
| | 5A10 | 97.7 | 4869.3 | Specimen_001_B5_B05_015.fcs |
| | 5D4 | 99.5 | 12933.3 | Specimen_001_B6_B06_016.fcs |
| | 5D12 | 98.8 | 10118 | Specimen_001_B7_B07_017.fcs |
| | 5 E2 | 99.1 | 9648.6 | Specimen_001_B8_B08_018.fcs |
| | 5 E5 | 89.7 | 3820.5 | Specimen_001_B9_B09_019.fcs |
| | 5F5 | 96.9 | 5485.8 | Specimen_001_B10_B10_020.fcs |
| | 5A7 | 98.6 | 9980.6 | Specimen_001_B11_B11_021.fcs |
| | 5F8 | 99.3 | 7276 | Specimen_001_B12_B12_022.fcs |
| | 4G6 | 98.1 | 10644.7 | Specimen_002_C1_C01_023.fcs |
| MC38 blank | stained | 62.8 | 3405.6 | Specimen_001_A11_A11_009.fcs |
| | unstained | 0.5 | 351.8 | Specimen_001_A12_A12_010.fcs |

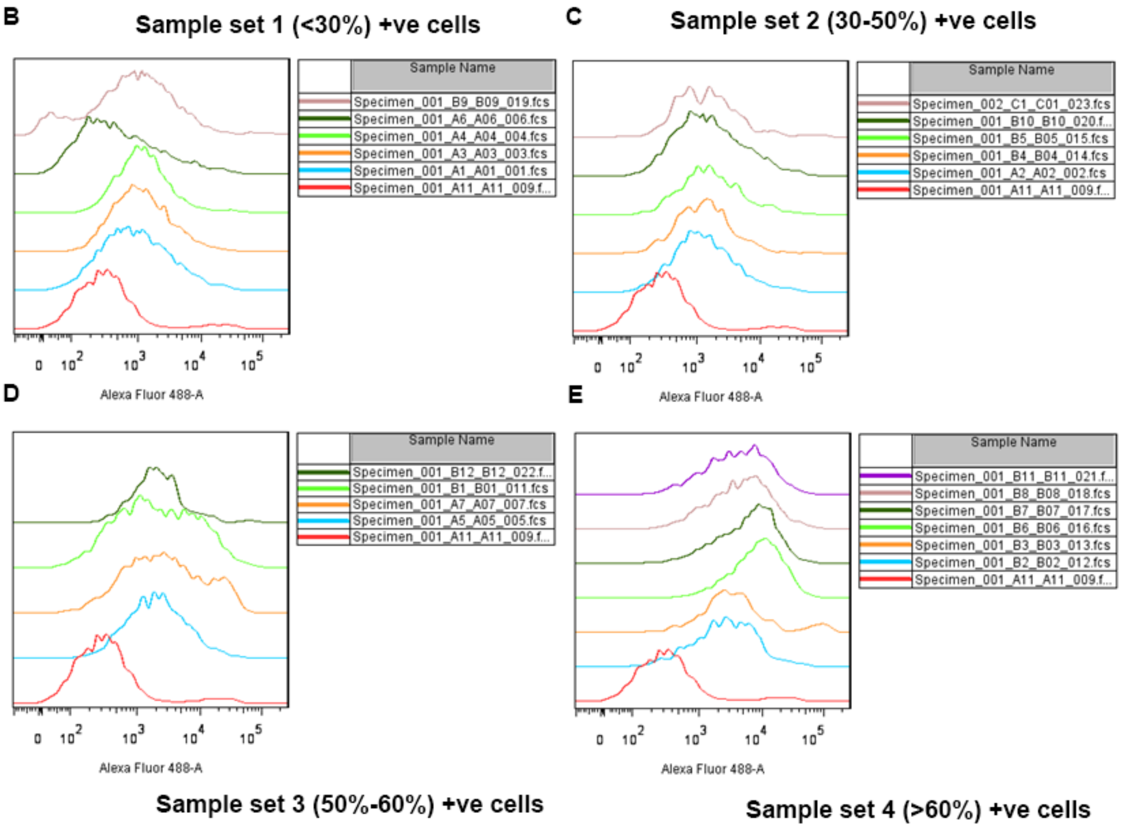


**Fig. S9. Validation of human PD-L2 over-expression in MC38 cells.** (A) Table showing percentage and mean fluorescent intensity of MC38 cells expressing 21 human PD-L2 individual clones. (B) MC38-hPDL2 cells showing less than 30% of hPD-L2 positive population. (C) MC38-hPD-L2 cells showing less 30%-50% of hPD-L2 positive population. (D) MC38-hPD-L2 cells showing 50%-60% hPD-L2 positive population. (E) MC38-hPD-L2 cells showing more than 60% hPD-L2 positive population.

**A**

| Clone | Alexa488 MFI | P2 %Parent | Peak number |
|---|---|---|---|
| Hep3B blank+Ab2 | 53.1 | 0.3 | 1 |
| Hep3B blank+Ab1+Ab2 | 2699 | 75.7 | 1 |
| 5C5 | 34409 | 98.4 | 1 |
| 5D8 | 53268 | 95.9 | 1 |
| 5E8 | 50584.3 | 95.3 | 1 |
| 5F8 | 32883.8 | 98.8 | 1 |
| 5F5 | 28832.1 | 98.9 | 1 |
| 4B9 | 41703.5 | 98.5 | 1 |
| 4G6 | 34652.9 | 98.1 | 1 |
| 4E2 | 38126.1 | 99.1 | 1 |
| 4F2 | 46515.5 | 92.7 | 1 |
| 4E7 | 32394.5 | 98.7 | 1 |
| 5E7 | 50304.4 | 98.6 | 1 |
| 5F7 | 39739.9 | 97.1 | 1 |
| 4F9 | 42523.6 | 96.4 | 1 |
| 5G3 | 43873.9 | 98.4 | 1 |
| 5G6 | 33266.8 | 98.5 | 1 |
| 5G9 | 61274.1 | 97.8 | 1 |

| Clone | Alexa488 MFI |
|---|---|
| 5G9 | 61274.1 |
| 5D8 | 53268 |
| 5E8 | 50584.3 |
| 5E7 | 50304.4 |
| 4F2 | 46515.5 |
| 5G3 | 43873.9 |
| 4F9 | 42523.6 |
| 4B9 | 41703.5 |
| 5F7 | 39739.9 |
| 4E2 | 38126.1 |
| 4G6 | 34652.9 |
| 5C5 | 34409 |
| 5G6 | 33266.8 |
| 5F8 | 32883.8 |
| 4E7 | 32394.5 |
| 5F5 | 28832.1 |

**B**

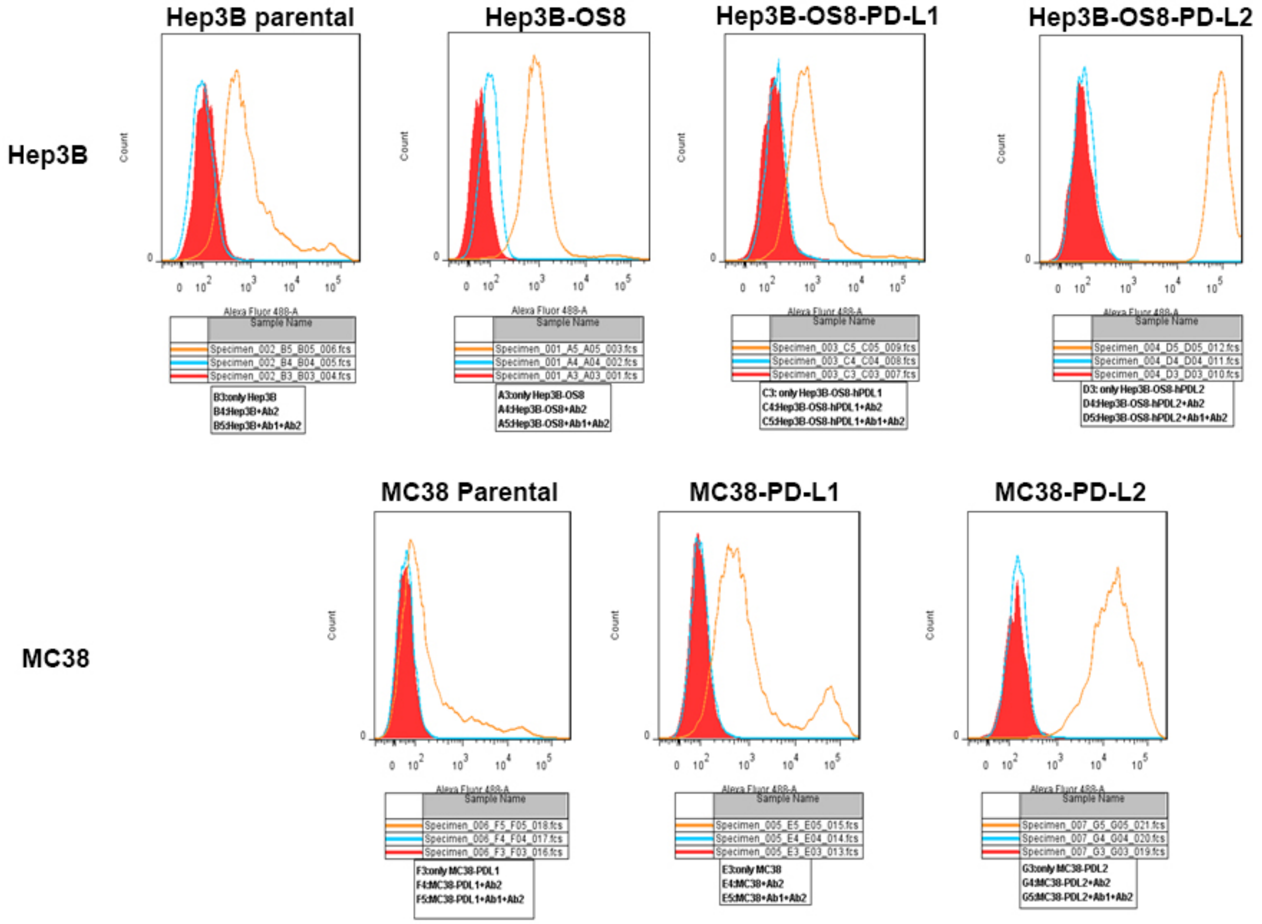


**Fig. S10. Validation of human PD-L2 expression on Hep3B-OS8 and MC38 cells** (A) Table showing mean fluorescent intensity and percent of Hep3B-OS8 cells transfected with 16 cDNA clones positive for human PD-L2 over-expression. (B-E) Positive clones were sorted from the lowest efficiency to highest efficiency on the right. Ab1 is anti-human PD-L2 antibody and Ab2 is control IgG.

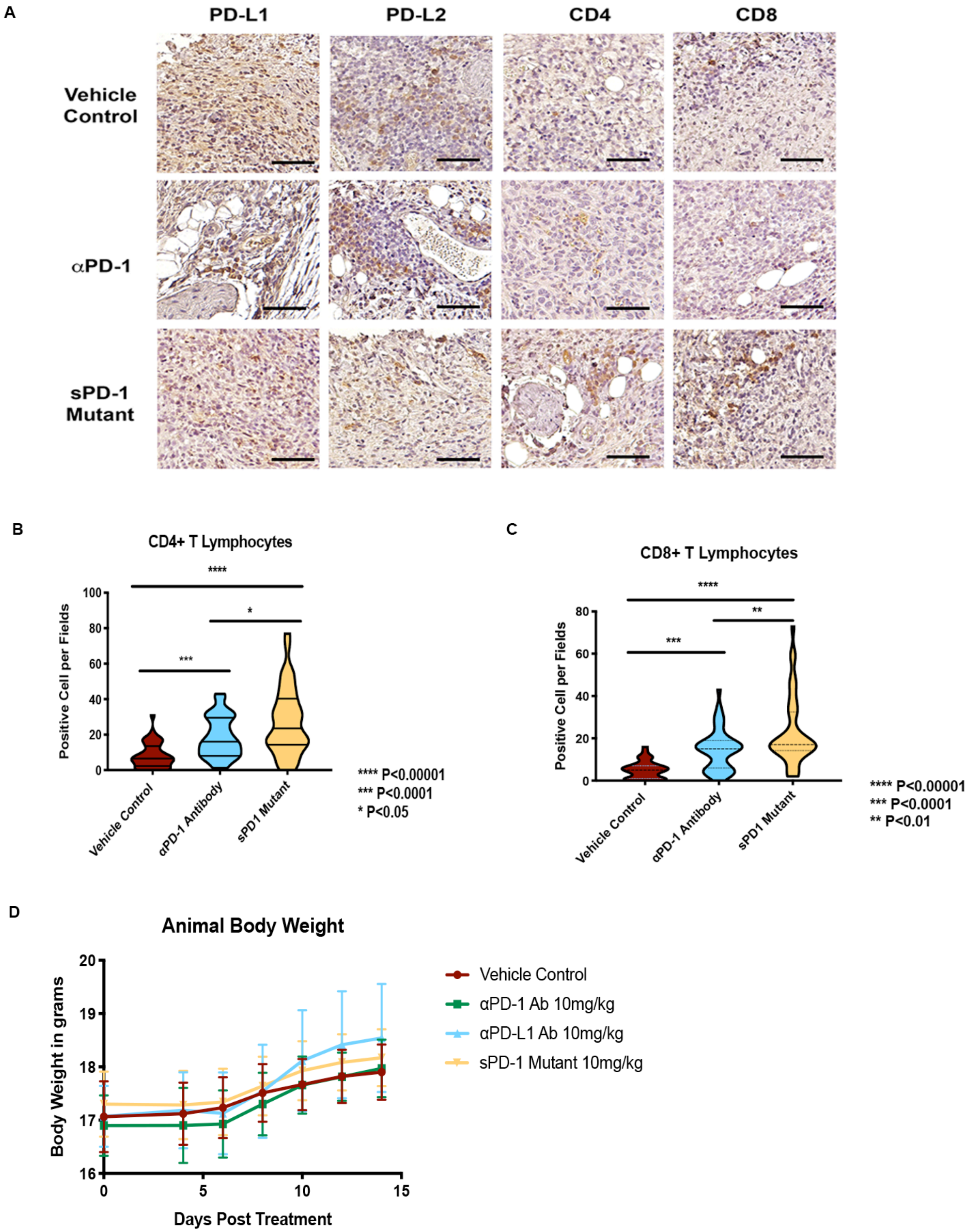
A
PD-L1
PD-L2
CD4
CD8
Vehicle Control
αPD-1
sPD-1 Mutant
B
CD4+ T Lymphocytes
Positive Cell per Fields
Vehicle Control
αPD-1 Antibody
sPD1 Mutant
**** P<0.00001
*** P<0.0001
* P<0.05
C
CD8+ T Lymphocytes
Positive Cell per Fields
Vehicle Control
αPD-1 Antibody
sPD1 Mutant
**** P<0.00001
*** P<0.0001
** P<0.01
D
Animal Body Weight
Body Weight in grams
Days Post Treatment
Vehicle Control
αPD-1 Ab 10mg/kg
αPD-L1 Ab 10mg/kg
sPD-1 Mutant 10mg/kg

**Fig. S11. sPD-1 mutant treatment leads to increased CD4+ and CD8+ TIL infiltration.** (A) The expression of PD-L1, PD-L2, CD4 and CD8 positive cells in ID8 tumors post-treatment was analyzed by immunohistochemical staining. Scale bar 30mm. (B) Violin quantitative plot of CD4+ T lymphocyte infiltrated into ID8 ovarian tumors post treatment. (C) Violin quantitative plot of CD8+ T lymphocyte infiltrated into ID8 ovarian tumors post treatment. (D) Mice body weight overtime during the UPK10 ovarian cancer study. Statistical analysis was conducted using One-way ANOVA for comparing between treatment groups and repeated ANOVA for changes occur over-time. P value *=<0.05, **=<0.01. ***=<0.001.

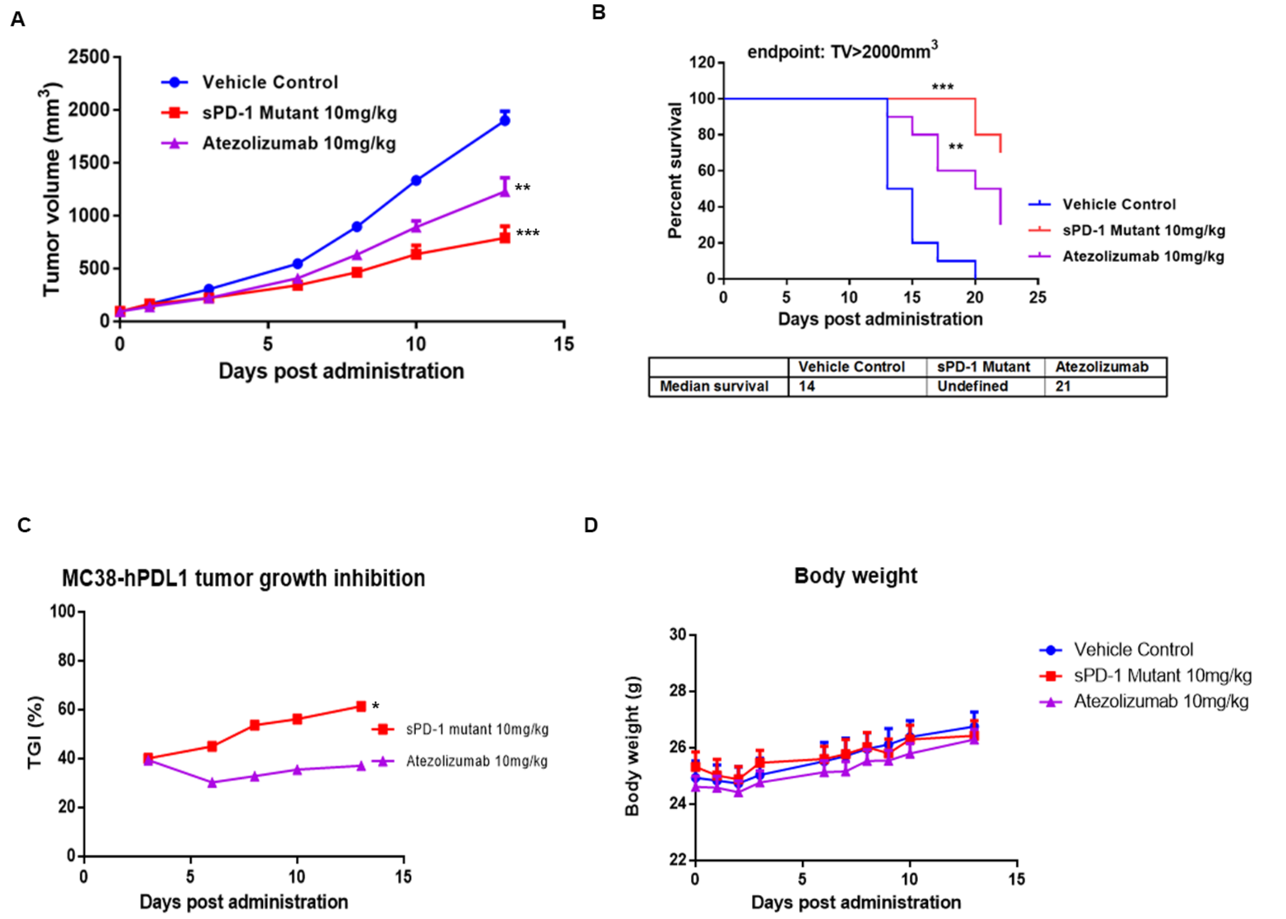


| | Vehicle Control | sPD-1 Mutant | Atezolizumab |
|---|---|---|---|
| Median survival | 14 | Undefined | 21 |

**Fig. S12. sPD-1 mutant treatment demonstrates superior anti-tumor activity in colorectal cancer compared to αPD-1 antibody** (A) C57B/6 mice inoculated with the MC39-hPD-L1 colorectal cancer then assigned to vehicle control, sPD-1 mutant 10mg/kg or Atezolizumab 10mg/kg. (B) Kaplan Meier survival plot showing animals been terminated upon reaching ethical endpoint for tumor growth with median survival listed below. (C) Percent of tumor growth inhibition comparing sPD-1 mutant and Atezolizumab treated groups. (D) Total body weight of mice treated with vehicle control, sPD-1 mutant and Atezolizumab during the experiment. N=10 for each treatment group. Error bar represents mean and standard deviation.

A

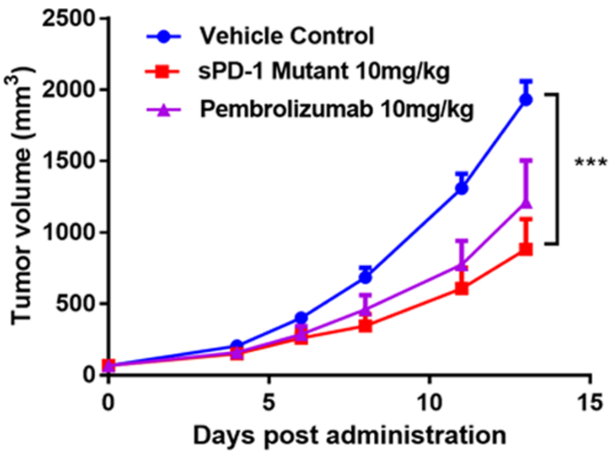


B

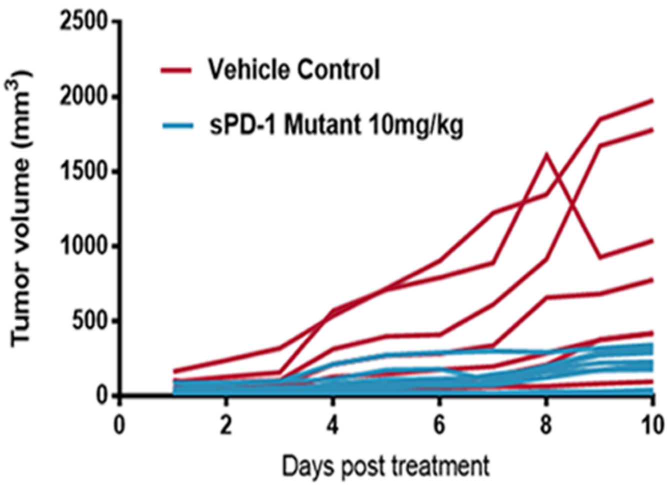


**Fig. S13. sPD-1 mutant treatment demonstrates superior anti-tumor activity in MC38 colorectal cancer over expressing PD-L2 and delays tumor growth in mouse melanoma model.** (A) Tumor growth over time in C57B/6 mice inoculated with MC38-hPD-L2 colorectal cancer then assigned to vehicle control, sPD-1 mutant 10mg/kg or Pembrolizumab10mg/kg. Each data point represents mean and SEM of individual tumor measured over-time. (B) Tumor growth over time in C57B/6 mice inoculated with B16/OVA melanoma cells then treated with to vehicle controls or sPD-1 mutant 10mg/kg.

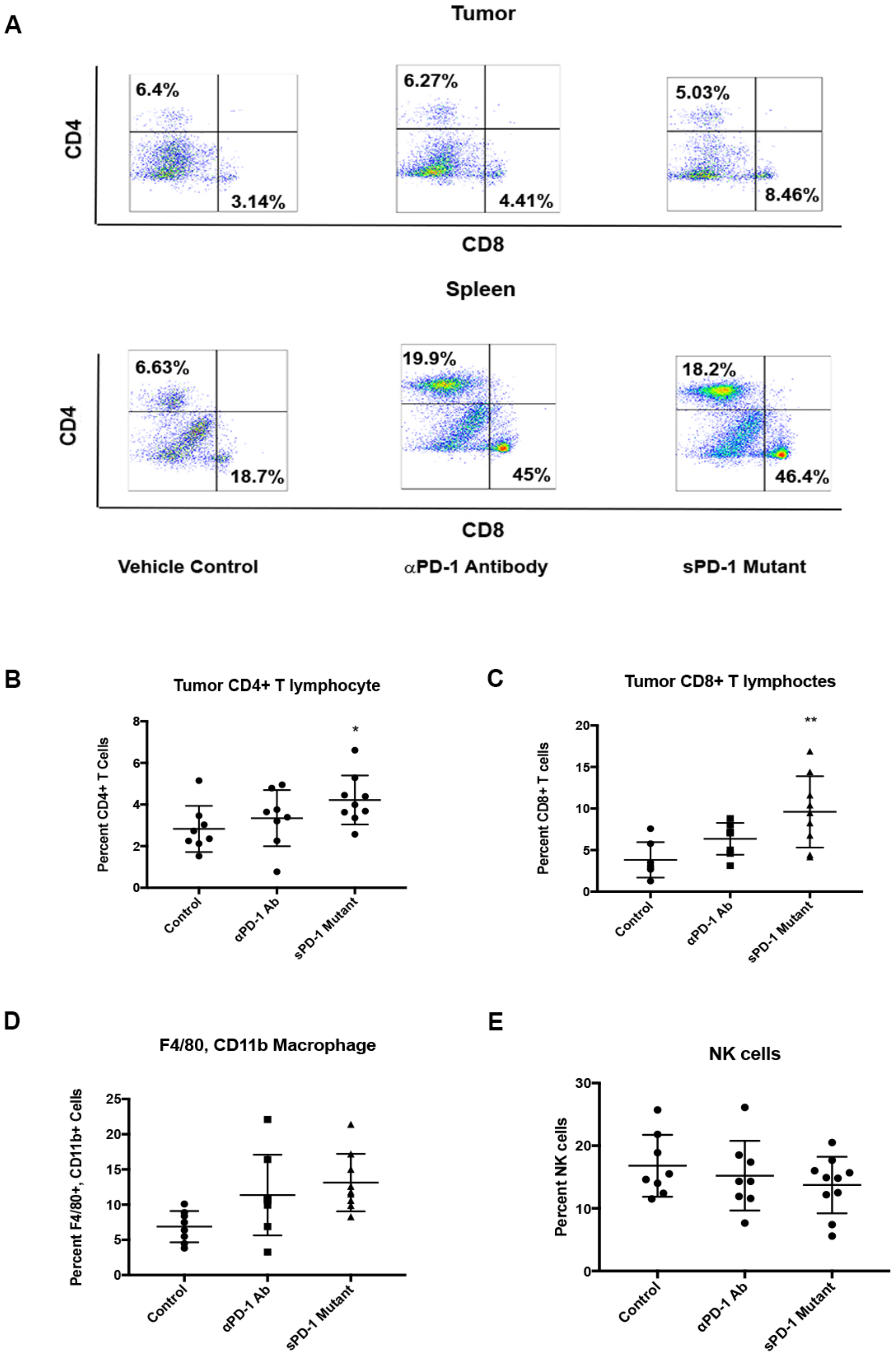

A
Tumor
6.4%
3.14%
6.27%
4.41%
5.03%
8.46%
CD4
CD8
Spleen
6.63%
18.7%
19.9%
45%
18.2%
46.4%
CD4
CD8
Vehicle Control
αPD-1 Antibody
sPD-1 Mutant
B
Tumor CD4+ T lymphocyte
Percent CD4+ T Cells
Control
αPD-1 Ab
sPD-1 Mutant
C
Tumor CD8+ T lymphoctes
Percent CD8+ T cells
Control
αPD-1 Ab
sPD-1 Mutant
D
F4/80, CD11b Macrophage
Percent F4/80+, CD11b+ Cells
Control
αPD-1 Ab
sPD-1 Mutant
E
NK cells
Percent NK cells
Control
αPD-1 Ab
sPD-1 Mutant

**Fig. S14. Immune profile of tumor associated immune cells in B16/OVA melanoma cells and MC38 tumor models treated with anti-mouse αPD-1 antibody and sPD-1 mutant.** (A) Representative flow cytometry dot plots of $CD4^+$ and $CD8^+$ cytotoxic T-cells isolated from tumor (upper panel) and spleen (lower panel) of B16/OVA melanoma tumors treated with vehicle control, anti-mouse aPD-1 blocking antibody 10mg/kg and sPD-1 mutant10mg/kg. (B) $CD4^+$ and (C) $CD8^+$ T cells isolated and analyzed from MC38 tumors treated with vehicle control, anti-mouse aPD-1 blocking antibody 10mg/kg and sPD-1 mutant 10mg/kg. (D) Percent positive NK cells in the tumors of MC38 tumors treated with vehicle control (N=8), anti-mouse αPD-1 antibody (N=8) and sPD-1 mutant antibody (N=10). Individual data point, mean and standard deviation shown. (B) Percent positive macrophages in the tumors of MC38 tumors treated with vehicle control (N=8), anti-mouse aPD-1 antibody (N=8) and sPD-1 mutant antibody (N=10). Individual data point, mean and standard deviation are shown. Statistical analysis was conducted using One-way ANOVA for comparing between treatment groups and repeated ANOVA for changes occur over time. P value *=<0.05, **=<0.01. ***=<0.001.

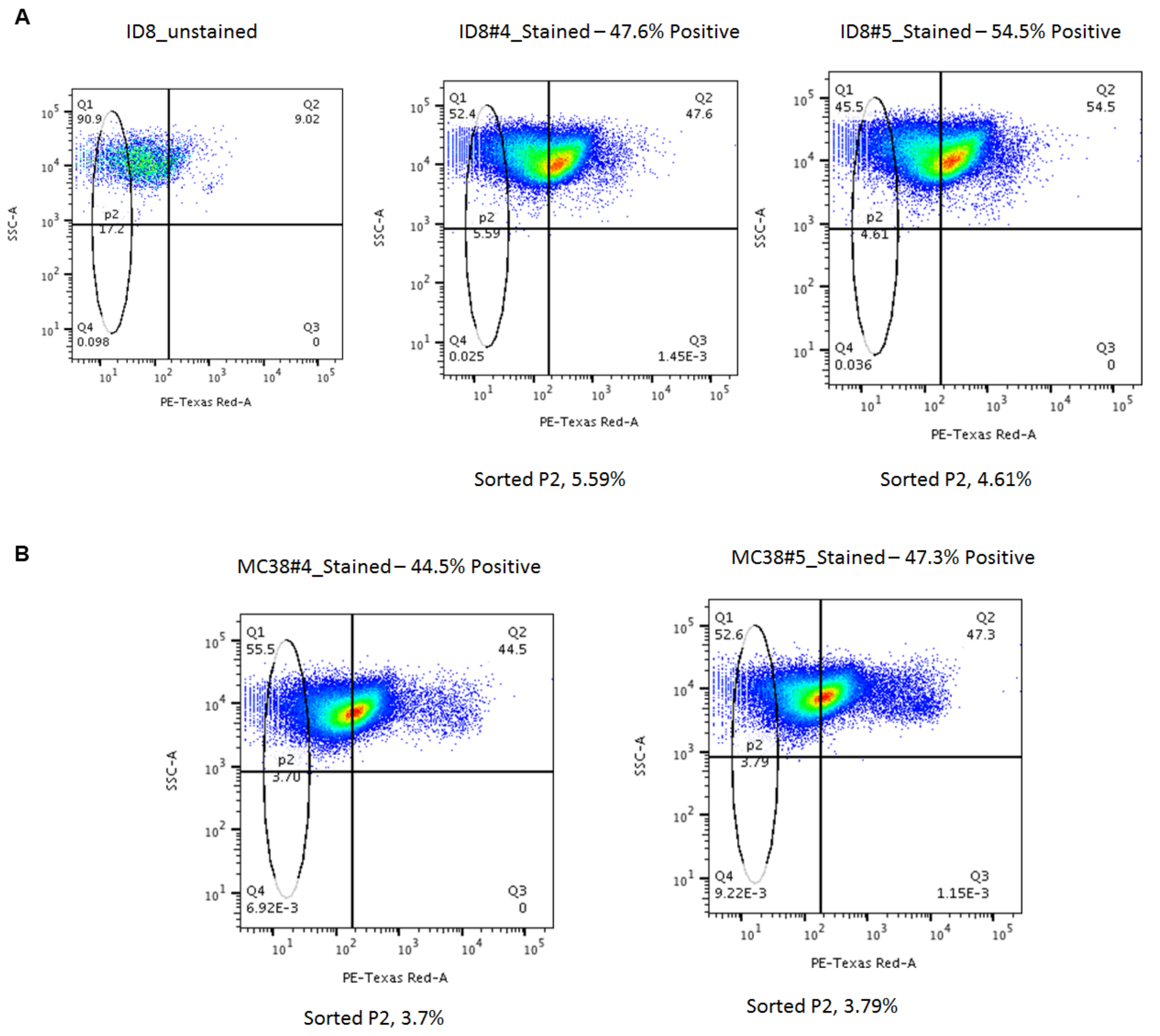


**Fig. S15. Isolation of PD-L1 negative cells post PD-L1 CRISPR transfection.** (A) PD-L1 negative ID8 cells sorted and collected post PD-L1 CRISPR clone 4 (left) and clone 5 (right) transfection. (B) PD-L1 negative MC38 cells sorted and collected post PD-L1 CRISPR clone 4 (left) and clone 5 (right) transfection.
