## Supplemental Table for "Neutralizing PD-L1 and PD-L2 Enhances the Efficacy of Immune Checkpoint Inhibitors in Ovarian Cancer"

### Supplemental Tables

**Table S1.** Sequencing of enriched sPD-1 library pool from each consecutive sort round. The residue number is listed at the top and the corresponding wild-type amino acid is given.

| | bp (Ig1) | AA (Ig1) | 33 | 34 | 36 | 37 | 45 | 49 | 50 | 57 | 59 | 70 | 74 | 76 | 77 | 84 | 87 | 88 | 89 | 91 | 92 | 93 | 96 | 99 | 101 | 102 | 103 | 107 | 110 | 112 | 113 | 114 | 116 | 119 | 124 | 125 | 127 | 131 | 132 | 135 | 139 | 140 | 145 | 146 | 147 | 148 |
|---|---|---|---|---|---|---|---|---|---|---|---|---|---|---|---|---|---|---|---|---|---|---|---|---|---|---|---|---|---|---|---|---|---|---|---|---|---|---|---|---|---|---|---|---|---|---|
| wt PD-1 | | | N | P | T | F | T | N | A | S | T | M | N | T | D | E | S | Q | P | Q | D | S | R | Q | P | N | G | H | V | R | A | R | N | G | G | A | S | K | A | K | R | A | T | E | R | R |
| S0.1 | 11 | 10 | | | | | | | | | A | | S | | | | G | | L | | | | | | | | | | | | | | | D | | V | | | V | | | V | | | K | G |
| S0.2 | 4 | 3 | D | | | | | | | | | | | | | | | | | | | | | | | | | | | | | | | | | | | | V | | G | | | | | |
| S0.3 | 7 | 5 | D | | | | | | | | A | | D | | | | G | | | | | | | | | | | | | | | | | | | | | | I | | | | | | | |
| S0.4 | 2 | 2 | | | | | | | | | A | | | | | | | | | | | | | | | | | | | | | | | | | | | | V | | | | | | | |
| S0.5 | --- | --- | | | | | | | | | | | | | | | | | | | | | | | | | | | | | | | | | | | | | | | | | | | | |
| S0.6 | 4 | 3 | | | | | | | | | | | D | | | | | | | | | R | | | | | | | | | | | | | | | | | V | | | | | | | |
| S0.7 | 6 | 4 | | | | | | | | | | | | | | | G | | L | | | | | | | | D | | | | | | | | | | | | V | | | | | | | |
| S0.8 | 9 | 9 | | S | | | | | | | | | D | | | | | | L | | | | | | | | | | | G | | | S | | S | | | | V | | | | | | K | G |
| S0.9 | 4 | 4 | | | I | | | | | | | | S | | | | | | L | | | | | | | | | | | | | | | | | | | | V | | | | | | | |
| S0.10 | 8 | 6 | | | | | | | | | A | | | | | | | | | | | | | | | | | | | | | | | | | | L | R | I | R | G | | | | | |
| S1.1 a | 10 | 6 | | | | | | | | | A | | | | | | G | | L | | | | | | | | | | | | | | | | | | | R | I | | | | A | | | |
| S1.1 b | 6 | 4 | | | | | | | | | | | | | | | | | L | | | | | | | | | | | | | | | | | | L | R | I | | | | | | | |
| S1.1 c | 9 | 7 | | | | | | | | | A | | | | | | | | | | | R | | | | | | | | | | | | | | | L | R | I | R | | V | | | | |
| S1.1 d | 3 | 3 | | | | | | | | | | | D | | | | | | | | | | | | | | | | | | | | | | | | | | V | | | V | | | | |
| S1.1 e | 6 | 5 | | | | | | | | | A | | | A | | | G | | | | | | | | | | | | | | | | | | | | | | I | | | V | | | | |
| S1.1 g | 10 | 5 | | | I | | | | | | | | | | | | | | | | | | | | | | | | | | | | | | | | F | | I | | | | | | K | G |
| S1.1 h | 5 | 4 | | | | | | | | | | | | | | | | | | | | | | | | | | | | | | | | | S | | L | | V | | | V | | | | |
| S1.1 i | 8 | 7 | | | | L | | | | | A | | | | | | G | | L | | | | | | | | | | | | | | S | | | | | | I | | | | I | | | |
| S1.2 a | 5 | 4 | | | | | | | | | | | | | | | G | | L | | | | | | | | | | | | | | | | | | | | I | | G | | | | | |
| S1.2 b | 6 | 4 | | | | | | | | | | | | | | | | | | | | | | | | | | | | | | | S | | | | L | R | I | | | | | | | |
| S1.2 c | 8 | 5 | | | | | | | | | | | | | | | | | | | | | | | | | | | | | | | | | | | L | R | I | R | G | | | | | |
| S1.2 d | 5 | 3 | | | | | | | | | | I | | | | | | | | | | | | | | | | | | | | | | | | | | | I | | G | | | | | |
| S1.2 e | 4 | 4 | | | | | | | | | | | | | | | G | | L | | | | | | | | | | | | | | | | | | | | V | | | V | | | | |
| S1.2 f | 11 | 7 | | | | | | | | | | | | | | | G | | | | | | | | | | | | | | | | | | | | L | R | I | R | | | | | | |
| S1.2 g | 9 | 8 | | | | | | | | | | | D | | | | G | | L | | | | | R | | | | | | | | | | | | | L | | V | | | V | I | | K | G |
| S1.2 i | 8 | 7 | D | | | | | | | | | | S | | | | | | | | | | | | | | | | | | | | | | | | L | | V | | | V | I | | K | G |
| S2.1 | 9 | 6 | | | | | | | | N | | | D | | | | G | | L | | | | | | | | | | | | | | | | | | | | I | | | V | | | | |
| S2.2 | 6 | 5 | | | | | | | | N | | | | | | | G | | L | | | | | | | | | | | | | | | | | | | | I | | | V | | | | |
| S2.3 | 8 | 6 | | | | | | | A | | A | | | | | | | | | | | | | | S | | | | | | | | | | | | L | | I | | | V | | | | |
| S2.4 | 6 | 5 | | | | | | | | | A | | | | | | | | | | | | | | | | | | | | | | | | S | | | | I | | | V | | K | | |
| S2.5 | 9 | 7 | | | | | | | | | | | | | | | G | | L | | | | | | | | | | | | | | | | S | | L | R | I | R | | | | | | |
| S2.6 | 10 | 8 | | | | | | | | | | | | | | | | | L | | | | | | | | | | | | | | T | | | | L | R | I | R | | V | I | | | |
| S2.7 | 7 | 5 | | | | | | | | | | | | | | | G | | L | | | | | | | | | | | | | | | | | | L | R | I | | | | | | | |
| S2.8 | 7 | 5 | | | | | | | | | | | D | | | | G | | L | | | | | | | | | | | | | | | | S | | | | I | | | | | | | |
| S2.9 | 9 | 8 | | | | | | | | | A | | | | | | G | | L | | | | | R | | | | | | | | | | | S | | | | V | R | | V | | | | |
| S2.10 | 9 | 7 | | | | | | | | | A | I | | | | | G | | L | | | | | | | | | | | | | | | | S | | | | V | | | V | | | | |
| S2.11 | 13 | 10 | | | | | | | | N | | | D | | | | | | | | | | | | | | | | | | | K | S | | | | L | R | I | R | | | | | K | G |
| S2.12 | 6 | 5 | | | | | | | | | A | | D | | | | G | | | | | | | | | | | | | | | | | | | | | | I | | | V | | | | |
| S2.13 | 9 | 8 | D | | | | | | | | | | | | | | G | | L | | | | | | | | | | | | | | | | | | L | R | I | R | | V | | | | |
| S2.14 | 12 | 8 | | | | | | | | | A | I | | | | | | | | | | | | | | | | | | | | | | | S | | L | | I | | G | | | | K | G |
| S2.15 | 8 | 7 | | | | | | | | | | I | | | | | G | | L | | | | | | | | | | | | | | S | | | | | | I | | | V | | | G | |

| | | | | | | | | | | | | | | | | | | | | | | | | | | | | | | | | | | | | | |
|---|---|---|---|---|---|---|---|---|---|---|---|---|---|---|---|---|---|---|---|---|---|---|---|---|---|---|---|---|---|---|---|---|---|---|---|---|---|
| S3.1 | 9 | 8 | | I | | | | A | | | | | | L | | | | G | | | | | | | S | S | | | | I | | | V | | | | |
| S3.2 | 3 | 2 | | | | | | | | | | | | | | | | | | | | | | | | | | | | V | | | V | | | | |
| S3.3 | 6 | 5 | | | | | | | | | | | G | L | | | | | | | | | | | | S | | | | I | | | V | | | | |
| S3.4 | 10 | 8 | | | | | | I | | D | | | G | L | | | | | | | | | | | S | S | | | | I | | | V | | | | |
| S3.5 | 8 | 7 | | | | | | | | D | | | G | L | | | | | | | | | | | | S | | | R | I | | | V | | | | |
| S3.6 | 15 | 13 | | | L | | | | | D | | | G | L | | | | | | | R | | | | S | S | | L | R | I | | G | | | | K | G |
| S3.7 | 15 | 9 | D | | | | | | | | | | | L | | | | | | | | | | | | S | | L | R | I | R | G | | I | | | |
| S3.8 | 13 | 10 | D | | | | | | | S | | | G | | | | | | | | | | | | S | | | L | R | I | | | | | | K | G |
| S3.9 | 12 | 10 | | | | | | | | D | | | G | L | | | | | | | | | G | | S | S | | | | V | | | V | | | K | G |
| S3.10 | 8 | 7 | | | | | | A | | D | | | G | L | | | | | | | | | | | | S | | | | I | | | V | | | | |
| S3.11 | 13 | 10 | | | | A | | | | | | | G | L | | S | | | R | | | | | V | | S | | | | I | R | G | | | | | |
| S3.12 | 6 | 5 | | | | | | | | | | | G | | | | | | | | | | | | D | S | | | | I | | | V | | | | |
| S3.13 | 6 | 5 | | | | | | | | D | | | | L | | | | | | | | | | | | S | | | | I | | | V | | | | |
| S3.14 | 9 | 7 | | | | A | | | | | | | G | L | | | | | | | | | G | | S | S | | | | I | | | | | | | |
| S3.15 | 3 | 2 | | | | | | | | | | | | | | | | | | | | | | | | | | | | V | | | V | | | | |
| S4.1 | 11 | 8 | | | | | | | | | | | G | L | | | | | | | | | | | | S | | L | | I | | V | | | | K | G |
| S4.2 | 8 | 7 | | | | A | | | | | | | G | L | | | | | | | | | | | T | S | | | | I | | | V | | | | |
| S4.3 | 11 | 8 | | | | | | | | | | | G | L | | | | | | | | | | | S | S | | L | R | I | R | | | | | | |
| S4.4 | 14 | 11 | | | | | | | | | | | G | L | | | | | | | | | | | S | S | | L | R | I | R | G | | | | K | G |
| S4.5 | 7 | 6 | | | | | | | | | | | G | L | | | | | | | | | | | S | S | V | | | I | | | | | | | |
| S4.6 | 8 | 7 | | | | | N | | | | | | G | L | | | | | | | | | | | S | S | | | | V | | | V | | | | |
| S4.7 | 7 | 6 | | | | | | | | | | | G | L | | | | | | | | | | | S | S | | | | I | | | V | | | | |
| S4.8 | 10 | 7 | | | | | | | | | | | G | L | | | | | | | | | | | | S | | L | | I | | | V | | K | | |
| S4.9 | 8 | 7 | | | | A | | | | | | | G | L | | | | | | | | | | | T | S | | | | I | | | V | | | | |
| S4.10 | 11 | 9 | | | | A | N | | | | | | G | | R | | | | | | | | | V | | S | | L | R | I | | | | | | | |
| S4.11 | 9 | 6 | | | | | | | | | | K | G | L | | | | | | | | | | | | S | | | | I | | | V | | | | |
| S4.12 | 7 | 6 | | | | | | | | | | | G | L | | | | | | | | | | | S | S | | | | I | | | V | | | | |
| S4.13 | 8 | 6 | | | | | | | | | | | | | | A | | | | | | | | | S | S | | L | R | I | | | | | | | |
| S4.14 | 13 | 10 | | | | A | | | | | | | G | L | | A | | | R | | | | | V | | D | | | | I | R | G | | | | | |
| S4.15 | 8 | 7 | | | | | | | | | | | G | L | | | | | | | | | | | S | S | | | R | I | | | V | | | | |
| S5.1 | 10 | 7 | | | | | | | | | | | G | L | | | | | | | | | | | S | S | | V | | I | | | V | | | | |
| S5.2 | 8 | 7 | | | | | | | | | | | G | | | | | | R | | | M | | | S | S | | | | I | | | V | | | | |
| S5.3 | 7 | 6 | | | | | | | | | | | G | | R | | | | R | | | | | | S | S | | | | I | | | | | | | |
| S5.4 | 13 | 10 | | | | | | | | | | | G | | R | | | | | | R | | | | S | S | | L | | I | | G | | | | K | G |
| S5.5 | 9 | 8 | | | | | N | | | | | | G | L | | G | | | | | | | | | | S | | | | I | | | V | | | K | |
| S5.7 | 9 | 7 | | | | | | | | | | | G | L | | | | | | | | | | | S | S | | L | | I | | | V | | | | |
| S5.9 | 13 | 11 | | | | | | | V | D | | | G | L | | | R | | | S | | | | | | S | | L | | I | | | V | | | | |
| S5.10 | 8 | 6 | | | | | | | | | | | G | | | G | | | | | | | | | S | S | | | | I | | | V | | | | |
| S5.12 | 12 | 10 | D | | | | | | | | N | | G | | | G | | G | R | | | | | | | S | | L | | V | | | V | | | | |

| | | | | | | | | | | | | | | | | | | | | | | | | | | | | | | |
|---|---|---|---|---|---|---|---|---|---|---|---|---|---|---|---|---|---|---|---|---|---|---|---|---|---|---|---|---|---|---|
| S6.8.1 | 13 | 11 | | | S | | | | | | G | L | | | | | R | | | S | S | L | R | I | R | G | | | | |
| S6.8.2 | 8 | 7 | | | | | | | | | G | | | | | R | | M | | S | S | | | I | | | V | | | |
| S6.8.3 | 12 | 10 | | | | | | | | | G | L | | A | | | | | | S | S | L | R | I | R | | V | | | |
| S6.8.6 | 18 | 15 | S | L | | V | | | | | G | L | | G | G | | | | | S | S | L | | I | R | G | | | K | G |
| S6.8.9 | 12 | 10 | | | S | | | | | | G | L | | | | | | | | S | S | L | R | I | | | V | A | | |
| S6.8.10 | 12 | 9 | | | | | N | | | | G | L | | | | R | | | | T | S | L | | I | | | V | | | |
| S6.8.12 | 12 | 10 | D | | | | | | | | G | L | | G | G | R | | | | | S | L | | I | | G | | I | | |
| S6.8.13 | 10 | 7 | | | | | | | | | G | L | | | | | | | | S | S | V | | I | | | V | | | |
| S6.12.1 | 12 | 9 | | | | | N | | | | G | L | | | | R | | | | T | S | L | | I | | | V | | | |
| S6.12.2 | 18 | 15 | S | L | | V | | | | | G | L | | G | G | | | | | S | S | L | | I | R | G | | | K | G |
| S6.12.3 | 10 | 8 | | | | | | | | | G | L | | | | | | | V | S | S | L | R | I | | | | | | |
| S6.12.4 | 9 | 8 | | | | | | | | | G | L | | G | G | R | | | | S | S | | | V | | | V | | | |
| S6.12.5 | 13 | 10 | | | | | | | | | G | | R | | | | R | | | S | S | L | | I | | G | | | K | G |
| S6.12.6 | 13 | 12 | | | | | | I | | | G | L | | | | | | | | S | S | L | R | I | R | G | V | | K | |
| S6.12.7 | 10 | 7 | | | | | | | | | G | L | | | | | | | | S | S | V | | I | | | V | | | |
| S6.12.8 | 12 | 10 | D | | | | | | | N | G | | | G | G | R | | | | | S | L | | V | | | V | | | |
| S6.12.9 | 10 | 8 | | | | | | | | | G | L | | | | | | | | S | S | V | R | I | | | V | | | |
| S6.12.13 | 13 | 11 | | | S | | | | | | G | L | | | | | R | | | S | S | L | R | I | R | G | | | | |

**Table S2.** Calculated changes in binding affinity and protein stability (in kcal/mol) between sPD-1 and PD-L1 with mutations within and outside of the binding interface.

| Mutations | d Affinity | d Stability (solvated) |
|---|---|---|
| 8 mutations | -19.03 | -69.92 |
| 7 mutations (N116S excluded ) | -20.08 | -66.99 |
| 4 outside mutations | -1.61 | -30.66 |
| 3 interface mutations | -24.78 | -30.58 |
| 2 mutations on missing loop | 1.31 | -12.08 |
